# Counter-regulation of RNA stability by UPF1 and TDP43

**DOI:** 10.1101/2024.01.31.578310

**Authors:** Nicolas Gomez, Caroline Hsieh, Xingli Li, Megan Dykstra, Jacob Waksmacki, Christopher Altheim, Yoel Bechar, Joseph Klim, Benjamin Zaepfel, Jeffrey Rothstein, Elizabeth EM Tank, Sami J Barmada

## Abstract

RNA quality control is crucial for proper regulation of gene expression. Disruption of nonsense mediated mRNA decay (NMD), the primary RNA decay pathway responsible for the degradation of transcripts containing premature termination codons (PTCs), can disrupt development and lead to multiple diseases in humans and other animals. Similarly, therapies targeting NMD may have applications in hematological, neoplastic and neurological disorders. As such, tools capable of accurately quantifying NMD status could be invaluable for investigations of disease pathogenesis and biomarker identification. Toward this end, we assemble, validate, and apply a next-generation sequencing approach (NMDq) for identifying and measuring the abundance of PTC-containing transcripts. After validating NMDq performance and confirming its utility for tracking RNA surveillance, we apply it to determine pathway activity in two neurodegenerative diseases, amyotrophic lateral sclerosis (ALS) and frontotemporal dementia (FTD) characterized by RNA misprocessing and abnormal RNA stability. Despite the genetic and pathologic evidence implicating dysfunctional RNA metabolism, and NMD in particular, in these conditions, we detected no significant differences in PTC-encoding transcripts in ALS models or disease. Contrary to expectations, overexpression of the master NMD regulator UPF1 had little effect on the clearance of transcripts with PTCs, but rather restored RNA homeostasis through differential use and decay of alternatively poly-adenylated isoforms. Together, these data suggest that canonical NMD is not a significant contributor to ALS/FTD pathogenesis, and that UPF1 promotes neuronal survival by regulating transcripts with abnormally long 3’UTRs.

## Introduction

Amyotrophic lateral sclerosis (ALS) is a devastating neurodegenerative disease characterized by the progressive loss of upper and lower motor neurons^1,2^. How paralysis manifests and the extent to which cognitive function is involved varies significantly between individuals^3^. Despite this clinical and genetic heterogeneity, 95% of individuals with ALS exhibit cytoplasmic accumulation of TDP43 (transactive response element DNA/RNA binding protein, 43 kDa) in affected neurons and glia^4,5^. TDP43 is a ubiquitously expressed splicing factor that is primarily localized to the nucleus in healthy cells^6,7^. Mutations in the gene encoding TDP43 (*TARDBP*) and several similar RNA-binding proteins (RBPs) (e.g. FUS, HNRNPA2B1, MATR3) cause familial ALS as well as the related, often comorbid disease frontotemporal dementia (FTD)^8–10^.

In cellular and animal model systems, TDP43 overexpression results not only in neurodegeneration but also in TDP43 cytosolic mislocalization and aggregation, recapitulating key pathologic changes seen in humans^11–18^. We and others determined that overexpression of the RNA helicase up-frameshift 1 (UPF1), and to a lesser extent its binding partner UPF2, mitigates toxicity in both *in vivo* and *in vitro* models of ALS/FTD associated with TDP43 and FUS accumulation^12,13,19^. More recently, a similar neuroprotective effect was observed in *Drosophila* and neuron models of familial ALS/FTD due to *C9ORF72* hexanucleotide repeat expansions^20–22^, raising the possibility that UPF1 could act broadly to prevent neurodegeneration in ALS and FTD.

UPF1 is an obligate component of nonsense-mediated decay (NMD), a vital RNA surveillance pathway responsible for degrading premature termination codon (PTC)-containing transcripts that would otherwise produce truncated and potentially deleterious proteins^23,24^. These data are consistent with the detection of several new NMD substrates associated with TDP43 dysfunction^25–30^, as well as abnormal RNA stability in cells derived from ALS/FTD patients^31^. Little is known, however, about NMD activity in ALS/FTD models or patients, and whether any detected deficiencies are selective for these disorders.

Despite the importance of NMD to both human health and disease, quantifying NMD in biological systems remains an area of contention. Previous strategies employing overexpression-based reporter systems are limited in both generalizability and tractability^32,33^, particularly *in vivo*. Endogenous biomarkers of NMD activity identified in next-generation sequencing (NGS)-based screens, on the other hand, have taken an inherently limited “gene-centric” view of NMD that fails to take into account the vulnerability of differentially spliced transcript isoforms for a single gene^21,34^. Consequently, deriving NMD biomarkers from differential gene expression may suffer from low sensitivity and specificity.

To overcome these and other limitations we present an analytical pipeline, NMDq, that leverages available tools (STAR, StringTie2, and IsoformSwitchAnalyzeR), to identify novel, assayable, and highly sensitive NMD substrates harboring PTCs. We then use both a global and candidate-based approach to characterize the status of NMD in ALS/FTD models and tissues. Finally, we explore alternative RNA surveillance pathways through which UPF1 achieves neuroprotection, providing the basis for novel disease-modifying therapeutics for patients with ALS and FTD.

## Results

### NMDq RNA-seq analytic pipeline leverages available bioinformatic methods to quantify global NMD burden

Prior methods for assaying the clearance of PTC-containing NMD substrates have been limited in either isoform specificity or model generalizability. To overcome these limitations and quantify NMD at the transcript level in a variety of model systems, we implemented a modified version of a widely used RNA-seq analytic pipeline (**Fig. 1a**). Paired-end reads were aligned to the hg39 genome assembly using the splice-aware aligner STAR^35^. A *de novo* transcriptome was assembled using StringTie2^36,37^ and the GENCODE v38 annotation as a reference. The dataset-specific *de novo* transcriptome was then used to quantify aligned reads. Both annotated and novel transcripts were assessed for PTCs using the R package IsoformSwitchAnalyzeR^38^, which implements DEXseq^39^ to perform differential expression analysis of quantified transcripts.

**Figure 1:**
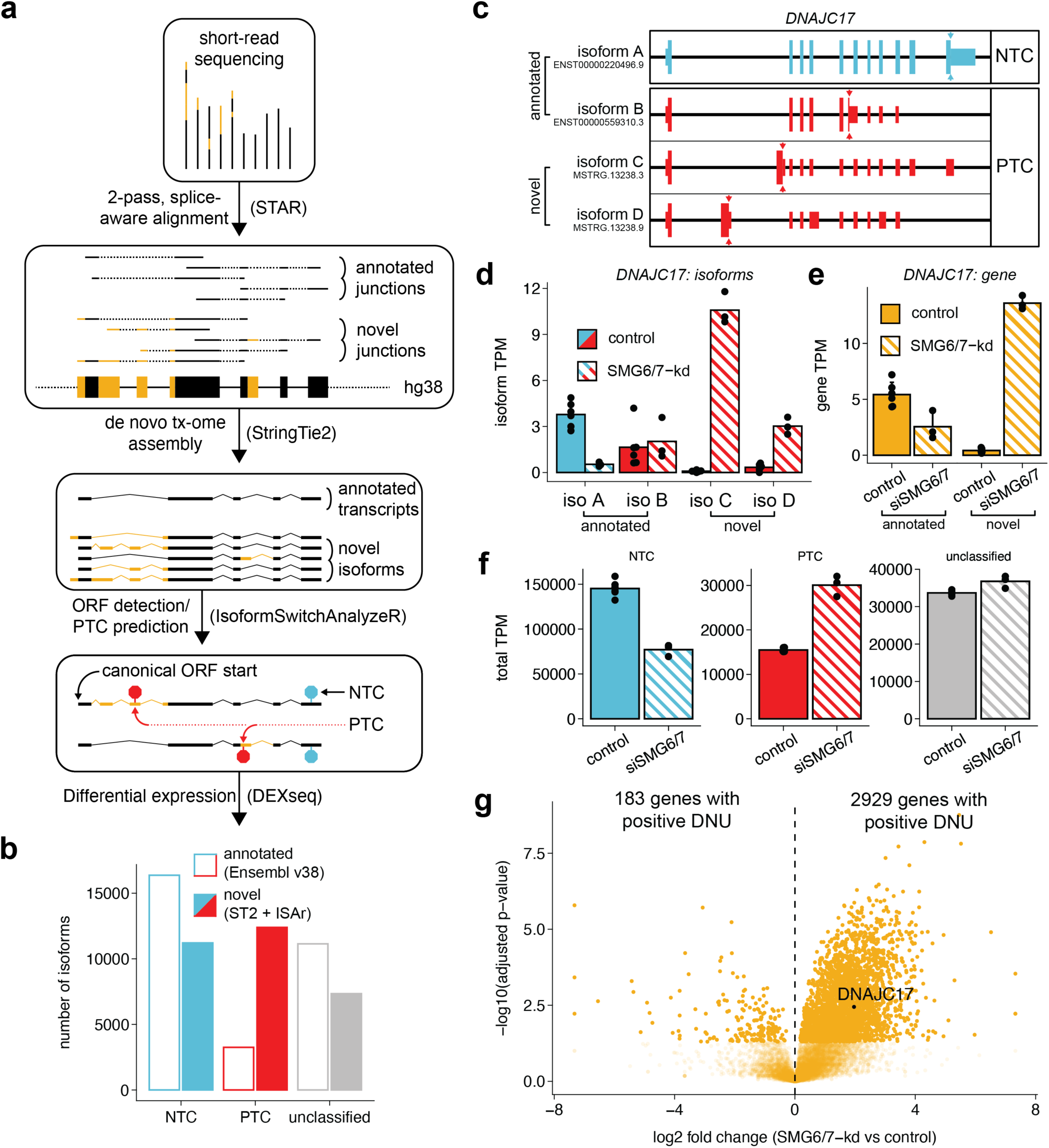
The NMDq analytic pipeline. **a,** Outline of NMDq raw data processing. **b**, Number of unique annotated and novel isoform models with normal termination codons (NTC), premature termination codons (PTC), and unclassified termination codons from the Colombo et al. (2017)^34^ dataset. **c**, Representative gene diagram for *DNAJC17,* and corresponding annotated and novel isoform models. **d**, Transcript abundance, measured as transcripts per million (TPM) for isoforms modeled in (**d**). **e**, gene-level TPM for DNAJC17. **f**, cumulative TPM of isoforms belonging to NTC, PTC, or unclassified categories. Data points in (**d**-**f**) represent biological replicates compared by Wilcoxon rank-sum test (p < 0.05, control n = 6, experimental n = 3). **g**, Volcano plot showing differential NMD-class usage (DNU). Data points represent log2 fold differences in PTC to total isoform abundance per gene between experimental and control groups. Abbreviations: STAR: Spliced Transcripts Alignment to a Reference; ORF = open reading frame; ST2: StringTie 2; ISAr: IsoformSwitchAnalyzeR; kd: knockdown.

As a proof of concept for the NMD quantification (NMDq) pipeline, we first applied it to a previous dataset wherein HeLa cells were treated with siRNA targeting essential NMD-pathway members SMG6 and SMG7, or non-targeting siRNA^34^. In each dataset, an approximately equal number of novel NTC- and PTC-containing isoforms were discovered through *de novo* transcriptome assembly, reducing the likelihood of bias towards either NMD class (**Fig. 1b**).

To determine the impact of unannotated gene products on measurements of NMD, we focused specifically on representative genes such as *DNAJC17* that display both NTC- and PTC-containing isoforms (**Fig. 1c-e**). Without the addition of novel isoforms, SMG6/7 knockdown appeared to selectively downregulate NMD-insensitive (i.e. NTC-containing) isoforms. In contrast, the inclusion of novel isoforms using NMDq demonstrated a pronounced effect of SMG6/7 knockdown on the abundance of NMD-sensitive (i.e. PTC-containing) isoforms, consistent with impaired surveillance of NMD substrates. To evaluate the cumulative consequence of NMD deficiency, isoform abundances were aggregated by NMD-class (**Fig. 1f**). Estimating the absolute NMD burden in this manner, we detected a >2-fold increase in PTC-containing isoforms upon SMG6/7 knockdown.

Alternative splicing and RNA-surveillance are stochastic processes capable of producing myriad low-abundance isoforms^40,41^. As a result, direct comparison of scarce NMD substrates (TPM<5) between groups can be particularly vulnerable to noise. To overcome this, isoforms were aggregated by NMD-class on a per-gene basis and used to determine differential NMD-class usage (DNU), representing the fraction of isoforms subject to NMD for each gene. DNU allows for direct comparison of the contribution of NMD to individual gene abundance, providing insight into those with similar sequence-dependent splicing potential in different biological contexts. In HeLa cells, DNU analysis identified 2929 genes with significantly increased NMD-class usage upon SMG6/7 knockdown, and 183 with increased NMD-class usage in the control group (log2 fold-change ≥ 1.5, adjusted p-value ≤ 0.05) (**Fig. 1g**). Together, these data confirm the utility of NMDq for measuring both global and gene-specific changes in the abundance of PTC-containing NMD substrates.

### NMDq is sensitive to both genetic- and pharmacologic-driven impairments in NMD in a variety of cell-based model systems

Due to the cell-specific nature of alternative splicing^42^ and NMD activity^43^, characterizing NMD in a universal, platform-independent manner can be challenging. Furthermore, because NMD involves the coordination of several distinct components, functional NMD inhibition can be achieved through several different mechanisms. To assess the sensitivity of NMDq across cell types and modes of NMD inhibition, we applied the pipeline to six distinct RNA-seq datasets (**Table 1**). NMDq was able to detect not just large changes, but also subtle differences in PTC-containing isoform abundances in HeLa cells, HEK293T cells, and human induced pluripotent stem cell (iPSC)-derived motor neurons (iMNs) when NMD was impaired by genetic or pharmacologic means (**Fig. 2**). Whereas differences in annotated isoforms alone suggested that the primary consequence of NMD inhibition was a loss of NTC-containing isoforms (**Fig. 2a-c, g-i**), the inclusion of novel unannotated isoforms demonstrated an unambiguous upregulation of PTC-containing transcripts in NMD-deficient conditions, consistent with direct inhibition of this pathway. In these experiments, UPF1 knockdown produced only a modest increase in NMD-substrates (**Fig. 2g**); however, measurement of NMD efficiency in this experiment could be complicated by difficulties in achieving adequate UPF1 knockdown, and the intrinsic toxicity of UPF1 loss of function^44^.

**Figure 2:**
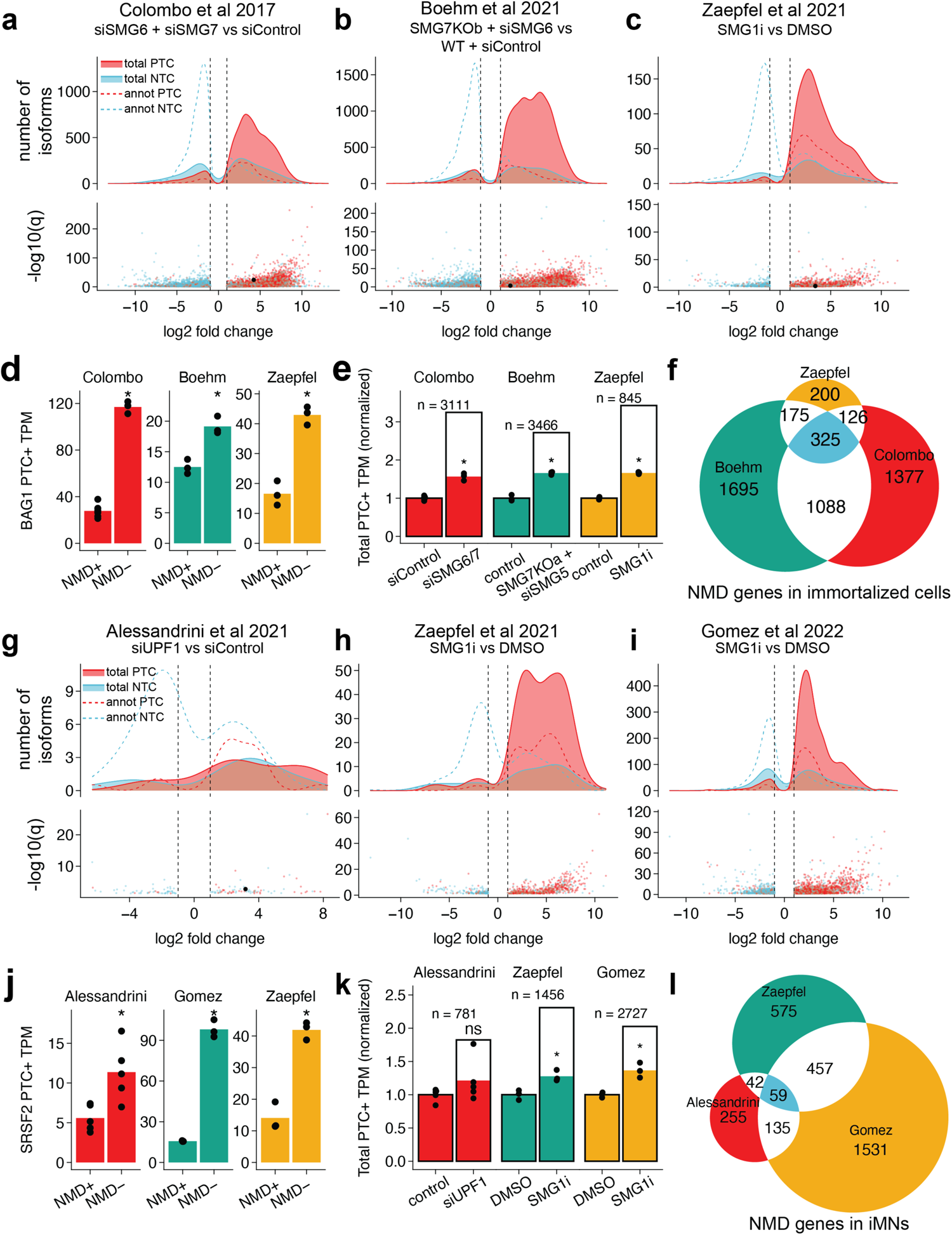
NMDq detects PTC burden in diverse cell types and conditions. **a**-**f**, Validation of NMDq in immortalized cells. **a**-**c**, **g**-**i**, Top, log2 fold-change density plots comparing relative isoform TPM between NMD-deficient (NMD-) and NMD-intact (NMD+) conditions. Bottom, volcano plots showing the distribution of significance. Dashed black lines indicate the filtering threshold of log2 fold-change ≤ −1 and ≥ 1. Black point corresponds to the commonest PTC-containing BAG1 isoform. All data shown are significant (adjusted p-value ≤ 0.05). **a**, siSMG6- and siSMG7-treated vs siControl-treated HeLa cells (from Colombo et al., 2017)^34^. **b**, siSMG6-treated SMG7-/- vs siControl SMG7+/+ HEK293T cells (from Boehm et al., 2021)^102^. **c**, SMG1 inhibitor (SMG1i)-treated vs DMSO-treated HEK293T cells (from Zaepfel et al., unpublished). **d**, **j**, cumulative, unnormalized TPM of PTC-containing BAG1 (**d**) or SRSF2 (**j**) isoforms in immortalized and iMN datasets, respectively. **e**, **k**, cumulative TPM belonging to PTC+ categories, normalized within the dataset to respective control. Data points represent biological replicates. Black outline represents the cumulative TPM of significantly differentially expressed isoforms (DNU+ genes). **f**, **l**, Euler diagram of the parental genes of differentially expressing PTC+ isoforms. **g**-**i**, Validation of NMDq in iMNs. **g**, siUPF1-treated vs siControl-treated cells (from Alessandrini et al., unpublished). **h**, SMG1i-treated vs DMSO-treated cells (from Zaepfel et al., 2021)^21^. **i**, SMG1i-treated vs DMSO-treated cells (this study).

**Table 1:**
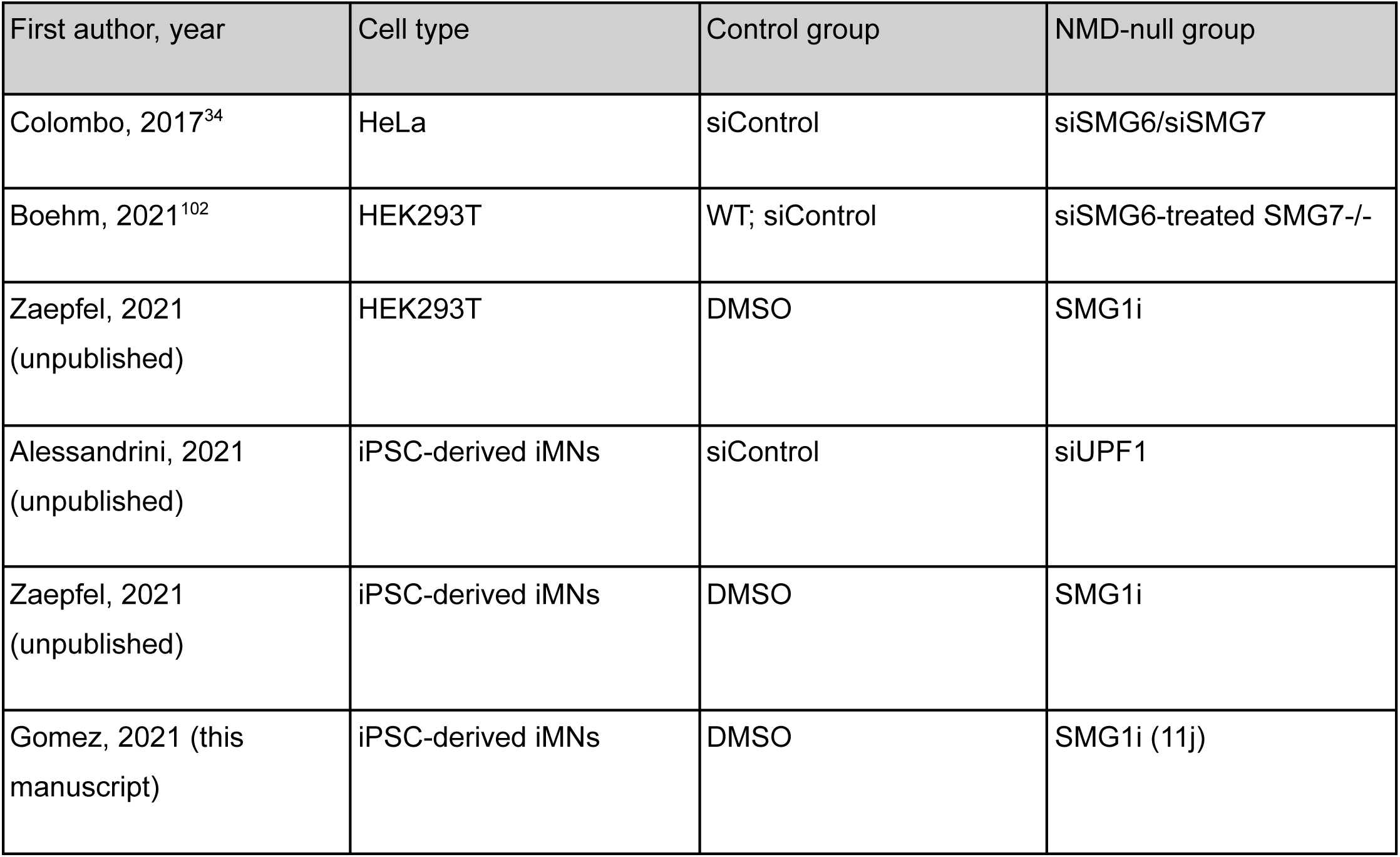
Validation RNA-seq datasets.

| First author, year | Cell type | Control group | NMD-null group |
| --- | --- | --- | --- |
| Colombo, 2017<sup>34</sup> | HeLa | siControl | siSMG6/siSMG7 |
| Boehm, 2021<sup>102</sup> | HEK293T | WT; siControl | siSMG6-treated SMG7<sup>-/-</sup> |
| Zaepfel, 2021<br>(unpublished) | HEK293T | DMSO | SMG1i |
| Alessandrini, 2021<br>(unpublished) | iPSC-derived iMNs | siControl | siUPF1 |
| Zaepfel, 2021<br>(unpublished) | iPSC-derived iMNs | DMSO | SMG1i |
| Gomez, 2021 (this manuscript) | iPSC-derived iMNs | DMSO | SMG1i (11j) |

Although DNU analysis can be helpful for gauging the contribution of NMD to the homeostasis of individual genes, we also sought to develop a singular measurement that encompasses NMD activity as a whole, enabling rapid comparison of NMD function across experimental conditions. We therefore summed the PTC-containing fraction of total reads and normalized this value to control, generating a global estimate of NMD burden (GNB). Total PTC+ abundance was greater in all NMD-deficient groups compared to respective controls (**Fig. 2g-h**, filled bars), consistent with effective inhibition of NMD in each condition. Because the majority of genes encode NMD-insensitive products (**Fig. 1b**), we performed a secondary, focused analysis of NMD substrates by limiting the PTC-containing fraction to intrinsically NMD-sensitive genes (INS), or those involved in statistically significant (q≤0.05) differential NMD-class switching (**Fig. 2g-j**, empty bars). The cumulative abundance of PTC+ isoforms encoded by INS genes was between 2 and 4-fold greater in NMD-deficient groups compared to respective controls. Moreover, many INS genes were common to at least two datasets: 325 genes were found in all immortalized cell datasets, and 59 of these were also detected within iMN datasets, suggesting the possibility of convergent and sensitive markers of NMD impairment. These candidates included several members of the serine-arginine rich splicing factor family, including *SRSF2*, *SRSF6*, *TRA2B* (also known as *SRSF10*). Each of these proteins regulates its expression by enhancing the inclusion of a PTC-containing “poison exon” within their cognate mRNA transcript, leading to its degradation by NMD^45^.

### NMDq identifies assayable targets of canonical NMD

To define universal markers of NMD impairment, we selected a series of novel PTC-containing NMD targets identified by NMDq that were expressed broadly, abundantly, and consistently across cell types. These conserved transcripts were compared by abundance within immortalized cells and iMNs (**Fig. 3a, Supp. Fig. 1a**). Nine highly abundant isoforms demonstrating prominent upregulation upon NMD inhibition were selected for RT-PCR-based validation (red points in **Fig. 3b, e**). HEK293T cells and iMNs were treated with either DMSO or the SMG1 inhibitor 11j^46^ for 48h and harvested for protein and RNA. The relative RNA abundance of selected PTC-containing isoforms were then compared between treatments. In keeping with the ability of NMDq to highlight conserved NMD substrates, all candidates showed significant upregulation upon treatment with 11j (**Fig. 3c, f**). Notably, novel NMD substrates identified by NMDq were considerably more sensitive as indicators of NMD impairment than previously published targets (*ATF3*, *ATF4*, and *ERN1*)^47^. *BAG1* (BAG family molecular chaperone regulator 1), a gene with a previously-identified NMD-sensitive transcript isoform^48^, showed a >12-fold increase upon 11j treatment in both HEK293T cells and iMNs. *SRSF2* exhibits highly-conserved poison exon sequences that are essential for the regulation of its expression via a process that requires NMD^45^. We observed a marked increase in poison exon usage in 11j-treated iMNs compared to DMSO-treated (representative examples given in **Supp. Fig. 1c**) suggesting that poison exon inclusion as a mode of NMD-mediated gene regulation is more widespread than previously appreciated.To determine if changes in transcript abundance upon NMD inhibition affect protein levels, we assayed one of the most abundant and broadly-expressed candidate NMD substrates, *SAT1*. In doing so, we observed a ∼1.5-fold increase in protein expression in 11j-treated HEK293T cells compared to controls (**Supp. Fig. 1b**), demonstrating that changes in NMD efficiency can have significant consequences at the protein level.

**Figure 3:**
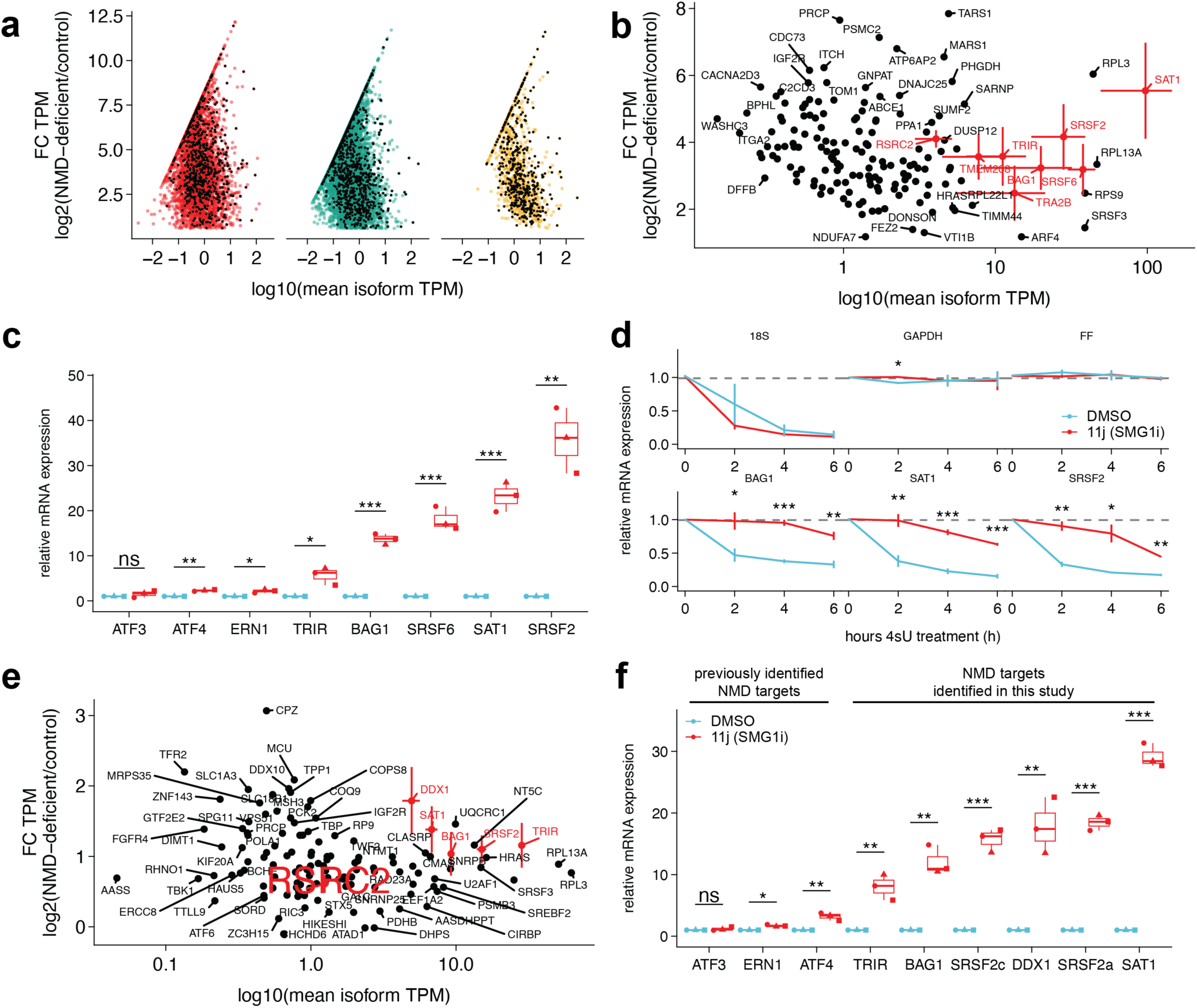
NMDq identifies biologically relevant and accessible targets of canonical NMD. **a**, First quadrant MA-plot (log-ratio vs mean expression) of immortalized cell validation datasets (from left to right: Colombo et al., 2017^34^: siSMG6- and siSMG7-treated vs siControl-treated HeLa cells; Boehm et al., 2021^102^: siSMG6-treated SMG7-/- vs siControl SMG7+/+ HEK293T cells; and Zaepfel et al., unpublished: SMG1 inhibitor (SMG1i)-treated vs DMSO-treated HEK293T cells). Points represent isoforms. Black points represent isoforms belonging to parental genes with at least one PTC-containing isoform up-regulated in all NMD-deficient conditions. **b**, **e**, the subset of isoforms in (**a**) and supplemental figure 1a that are significantly differentially expressed in all (**b**) or at least two datasets (**e**). Points in red show isoform candidates selected for follow-up studies. **c**, **f**, Isoform-specific RT-PCR of SMG1i- or DMSO-treated HEK293T cells (**c**) or iMNs (**g**). **d**, isoform-specific roadblock RT-PCR of SMG1i- or DMSO-treated HEK293T cells. DMSO = dimethylsulfoxide; FC = fold change; FF = firefly luciferase.

NMDq and all RNA-seq-based assays measure the steady-state abundance of transcripts and are therefore limited in their ability to accurately estimate the activity of NMD or other pathways that selectively degrade RNA. To determine whether the observed changes in the abundance of candidate NMD substrates were due to RNA turnover, we turned to roadblock-qPCR, a technique enabling the direct measurement of RNA clearance^49^. As expected, *GAPDH* and exogenous firefly luciferase RNA were stable and unaffected by NMD inhibition (**Fig. 3d**), while polyadenylated 18S ribosomal RNA was highly unstable^50^, and degraded rapidly through an NMD-independent process. Importantly, NMD-substrates identified by NMDq (*BAG1*, *SAT1*, and *SRSF2*) were unstable in DMSO-treated cells, and each transcript was significantly stabilized upon NMD inhibition, confirming their degradation through NMD.

### Global NMD burden is not consistently elevated in ALS models and human tissue

Although RNA metabolism is significantly altered in ALS/FTD^31^ patient samples, it remains unclear whether abnormalities in canonical NMD are responsible for these differences. Furthermore, overexpression of UPF1—an essential NMD factor—rescues phenotypes of disease in several ALS/FTD model systems^12,13,19–22^. These observations raise the possibility that NMD could be fundamentally impaired in ALS/FTD, and that UPF1 overexpression relieves this impairment. To address this question, we used NMDq to assess PTC-containing transcripts in several iMN models of ALS/FTD, including disease due to the *C9ORF72* hexanucleotide repeat expansion^51,52^, *TARDBP* mutation (M337V)^8,53^, TDP43 overexpression or knockdown^12^, and sporadic onset ALS (sALS) (**Table 2**). *C9ORF72* iMNs showed a subtle and nonsignificant decrease in GNB compared to control iMNs (**Fig. 4a**). These observations are consistent with previous studies suggesting that pathogenic *C9ORF72* mutations may in fact stimulate NMD activity^21,22^. iMNs derived from sALS iPSCs also showed a modest trend towards lower GNB compared to control iMNs. Conversely, TDP43 overexpression and knockdown, in addition to the M337V *TARDBP* mutation, all elicited mild and nonsignificant increases in GNB compared to controls (**Fig. 4b**). The impact of TDP43 overexpression on NMD activity was neither dose-dependent nor mutation-related: high-level TDP43 expression via lentiviral transduction had the same effect as a single copy insertion of TDP43, and overexpression of TDP43(M337V) was indistinguishable from TDP43(WT). Together, these data argue against NMD downregulation in patient-derived neuronal models of ALS/FTD.

**Figure 4:**
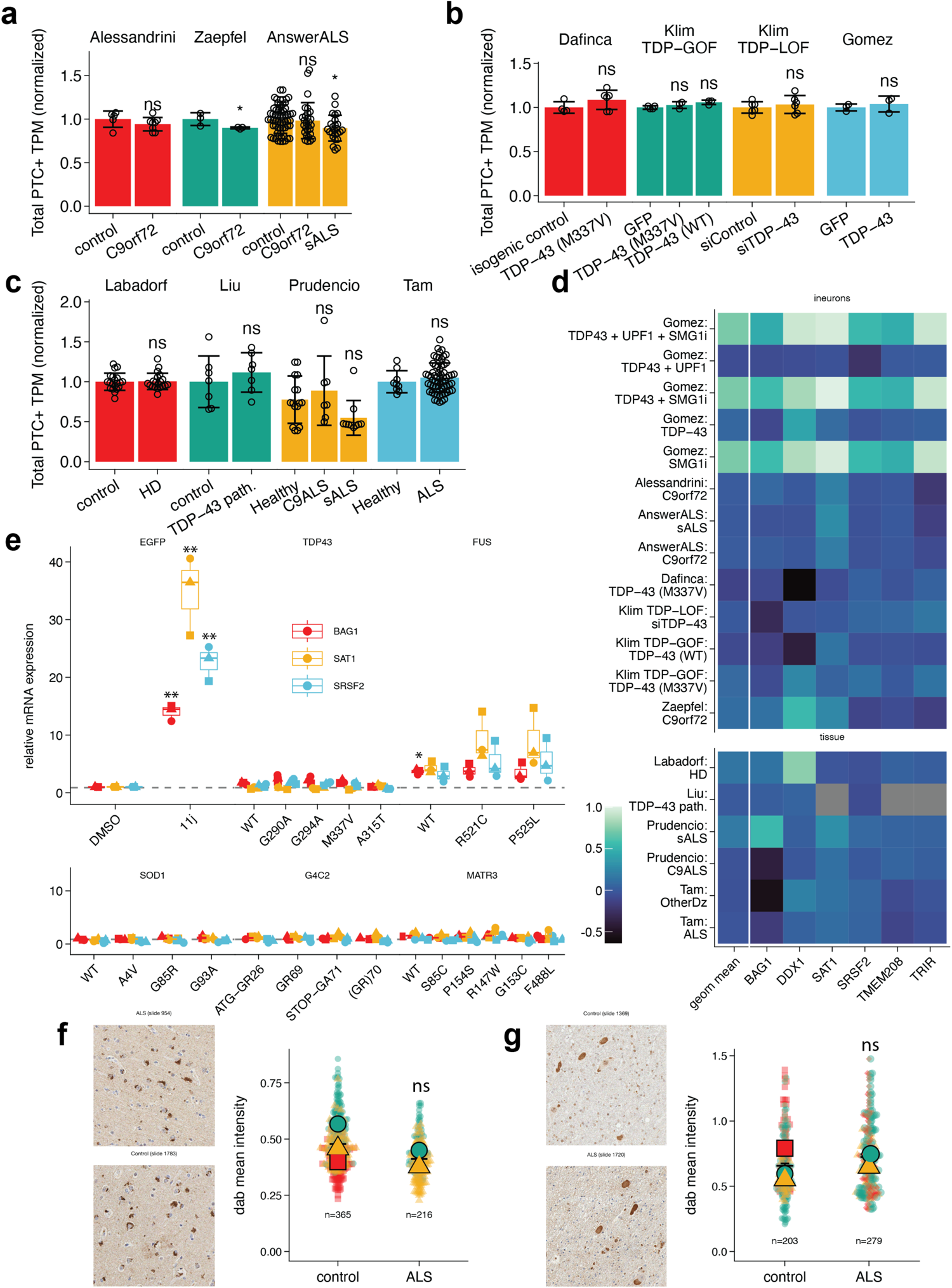
ALS models and tissues show no evidence for NMD impairment. **a**-**c**, Cumulative TPM belonging to the PTC category, normalized to experiment controls in *C9ORF72*- or sALS-based iMN models (**a**), TDP43-based iMN models (**b**), and post-mortem brain tissue (**c**). Data points represent biological replicates. Error bars represent standard error of the mean. Groups compared by Wilcoxon rank-sum test (*: p<=0.05). **d**, Control-normalized PTC+ fraction of NMD-sensitive targets identified by NMDq from RNA-seq datasets represented in **a**-**c**. Leftmost column represents the geometric mean of targets. **e**, qRT-PCR of NMD-sensitive isoforms in HEK293T overexpressing ALS-associated protein variants. Data points represent biological replicates from 3 experiments compared to the EGFP-DMSO condition by t-test. (*: p<=0.05, **: p<=0.01). **f**-**g**, phospho-UPF1 immunostaining of human spinal cord (**f**) and frontal cortex (**g**). Ghosted points represent cells from at least 2 individuals. Solid points represent means compared by Wilcoxon rank-sum test.

**Table 2:** ALS RNA-seq datasets.

| First author, year | Cell type | Control group | Experimental group | Disease Category |
| --- | --- | --- | --- | --- |
| Alessandrini, 2021<br>(unpublished) | iPSC-derived iMNs | isogenic control | *C9ORF72* mutation | *C9ORF72*-ALS/FTD |
| Zaepfel, 2021<br>(unpublished) | iPSC-derived iMNs | isogenic control | *C9ORF72* mutation | *C9ORF72*-ALS/FTD |
| AnswerALS, 2022<sup>103</sup> | iPSC-derived iMNs | isogenic control | *C9ORF72*-ALS/FTD and sALS | *C9ORF72*-ALS/FTD and sALS |
| Dafinca, 2020<sup>104</sup> | iPSC-derived iMNs | isogenic control | TDP43 (M337V) knock-in | *TARDBP*-ALS/FTD and sALS |
| Klim, 2019<sup>26</sup> | iPSC-derived iMNs | shControl or EGFP overexpression | TDP43 KD, TDP43 (WT) overexpression, TDP43 (M337V) overexpression | *TARDBP*-ALS/FTD and sALS |
| Liu, 2019<sup>65</sup> | FACS-sorted nuclei from patient cortex | TDP43 positive (no nuclear clearance) | TDP43 negative (nuclear clearance) | *C9ORF72*-ALS/FTD |
| Prudencio, 2015<sup>75</sup> | patient frontal cortex | Healthy control cortex | C9ALS or sALS | *C9ORF72*-ALS/FTD and sALS |
| Tam, 2019<sup>105</sup> | patient cortex | Healthy control cortex | ALS | *C9ORF72*-ALS/FTD and sALS |
| Labadorf, 2015<sup>54</sup> | patient prefrontal cortex | Healthy control cortex | Huntington's disease cortex | Huntington's disease |

Analogous trends were observed in human tissues (**Fig. 4c**). Using data from three separate studies of *C9ORF72*-linked ALS, we noted no significant change in the GNB to suggest changes in overall NMD activity. In sALS, we detected only a modest reduction in GNB, similar to what we originally found in sALS iMNs (**Fig. 4b**). A separate analysis of RNA-seq data from Huntington’s disease brain^54^ also showed no substantial differences from controls. Due to variability in sample preparation and sequencing methods across ALS datasets, we sought to validate these findings by specifically focusing on the abundance of NMDq-derived NMD-substrates within each of the iMN and human tissue datasets described in **Fig. 4a-c**. In each case, we observed little to no change in NMD substrates (**Fig. 4d**). As a positive control, we detected a robust increase in the abundance of the same set of NMD substrates in iMNs after the application of the NMD inhibitor 11j. These results suggest that NMD is not significantly affected in ALS/FTD or iMN models of these diseases.

The above studies are limited by data availability and the scarcity of samples from individuals with familial ALS/FD. Therefore, to examine the possibility that rare pathogenic mutations associated with ALS impact NMD, we expressed a broad array of ALS/FTD-associated proteins bearing disease-associated mutations in HEK293T cells, and measured the abundance of three NMD candidate substrates identified by NMDq (*BAG1*, *SAT1*, and *SRSF2*) by RT-qPCR (**Fig. 4e**). As expected, application of 11j resulted in a marked increase in all three candidates. However, among a panel of TDP43, FUS, SOD1, C9ORF72, and MATR3 variants, only FUS overexpression resulted in significant upregulation of NMD substrates. These results are consistent with previous studies suggesting that pathogenic FUS variants impair NMD. Even so, the majority of pathogenic variants displayed little to no effect on the same NMD substrates, arguing against a significant impact of TDP43, SOD1, C9ORF72, and MATR3 variants on NMD activity.

### UPF1 and TDP43 counter-regulate the essential arginine biosynthetic enzyme ASS1

In the absence of detectable NMD impairment in ALS disease models, we broadened our search for molecular pathways responsible for UPF1-mediated neuroprotection. We reanalyzed our lentivirus-treated iMN RNA-seq data, focusing on genes showing significant changes upon TDP43 overexpression that were subsequently rescued by co-expression of UPF1. Several candidates displaying counter regulation by TDP43 and UPF1 emerged from this analysis, including transcripts encoding ly6/PLAUR domain-containing protein 1 (LYPD1), laminin subunit alpha-4 (LAMA4), and argininosuccinate synthetase 1(ASS1) (**Fig. 5a**). LYPD1 is a neurotransmitter receptor binding protein of unknown function that is enriched in Von Economo neurons^55^ and layer 5b cortical neurons that are selectively vulnerable in FTD^56,57^ and mouse models of SOD1 ALS^58^, respectively. LAMA4 is an extracellular matrix protein of particular importance to neuromuscular junction physiology^59^, while ASS1 is an essential enzyme that catalyzes the rate-limiting step in arginine biosynthesis. Mutations affecting ASS1 function or expression cause familial citrulinemia, a pediatric neurological disease characterized by seizure and delayed cognitive development^60^. Notably, rescue by UPF1 was unaffected by 11j for each of these candidates, suggesting a mechanism independent of UPF1 phosphorylation.

**Figure 5:**
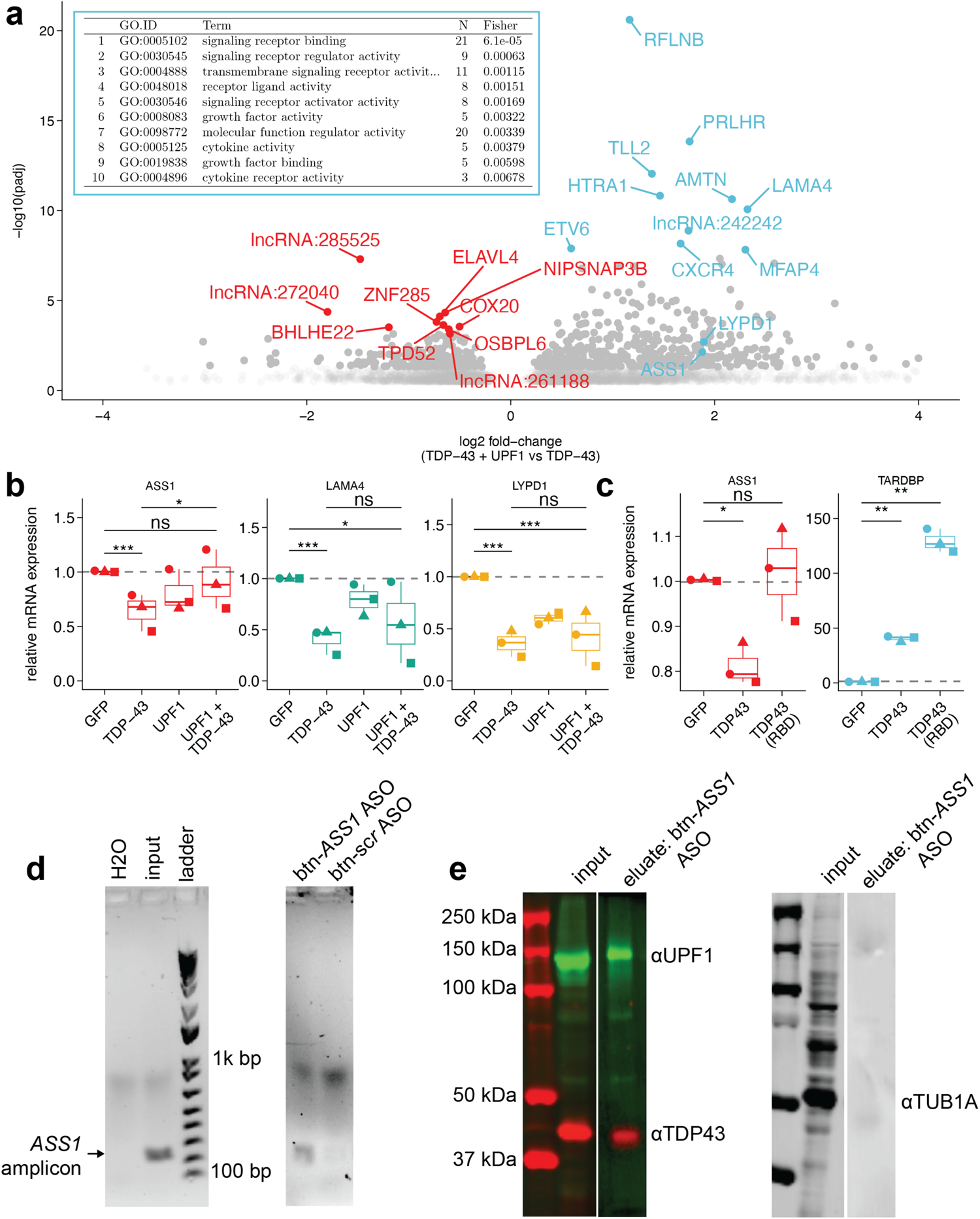
UPF1 counter-regulates TDP43-associated genes. **a**, Volcano plot of TDP43 + GFP vs TDP43 + UPF1 lentivirus-treated iMNs. **a, inset**, top 10 GO terms of significantly upregulated genes in this comparison. **b**, iMN qRT-PCR validation of candidates identified via RNA-seq. **c**, qRT-PCR of *ASS1* and *TARDBP* HEK293T cells transfected with GFP, TDP43, or RNA binding-deficient TDP43. **b**-**c**, data points represent biological replicates for 3 experiments. Indicated comparisons made by Wilcoxon rank-sum test. **d**, Agarose electrophoresis of *ASS1* PCR products purified by biotinylated-ASO pulldown and cDNA reverse transcription. **e**, SDS-PAGE and immunoblotting of mRNP species purified by biotinylated-ASO pulldown, showing successful pulldown of UPF1 and TDP43, but not the negative control (TUB1A).

Due to its established role in neurologic disease, we pursued *ASS1* as a candidate indicator of UPF1’s effects. First, we verified the reciprocal effect of TDP43 and UPF1 on *ASS1* mRNA by RT-qPCR in human iMNs (**Fig. 5b**). To determine if TDP43 directly regulated *ASS1*, we transiently transfected HEK293T cells with either WT or an RNA-binding-deficient (RBD-null) TDP43 variant carrying the F147L and F149L mutations^61,62^. As in iMNs (**Fig. 4a, b**), *ASS1* was downregulated by TDP43 overexpression; this effect required RNA binding, as *ASS1* expression was unperturbed in TDP43 RBD-null expressing cells (**Fig. 5c**). These data suggest that TDP43 directly binds *ASS1* mRNA, consistent with the existence of multiple UG-rich repeats within the ASS1 transcript^63^. To confirm this possibility, we performed a streptavidin pull-down assay of *ASS1* RNA-containing ribonucleoprotein (RNP) particles^64^. UV-crosslinked HEK293T cells were lysed and incubated with biotin-conjugated antisense oligonucleotides (btn-ASO) which were then immobilized and washed on streptavidin-bound iron nanoparticles. We designed btn-ASOs to hybridize with either *ASS1* and confirmed the specificity of these probes via PCR of the target, *ASS1* mRNA (**Fig. 5d**), compared to a scrambled ASO control. We also assessed the composition of ASS1-containing RNPs by SDS-PAGE and Western blotting. *ASS1* btn-ASO eluates contained both TDP43 and UPF1 protein but lacked TUB1A, confirming assay specificity (**Fig. 5e**), and suggesting that *ASS1* mRNA interacts directly with both TDP43 and UPF1.

Cryptic splicing in the absence of functional TDP43 results in downregulation of essential factors such as stathmin-2 (STMN2) in ALS/FTD tissue^26,27^. To determine if *ASS1* is similarly dysregulated by TDP43 *in vivo*, and to validate *ASS1* levels as a potential biomarker of TDP43 activity, we utilized single-molecule fluorescence *in situ* hybridization (smFISH) to examine *ASS1* transcripts in post-mortem spinal cord from individuals with ALS (**Fig. 6a-c**). *STMN2* and *ASS1* transcripts were abundant in spinal neurons from the anterior horn of controls, but both transcripts were effectively depleted from ALS spinal neurons. Thus, similar to *STMN2*, *ASS1* transcripts are downregulated in ALS spinal neurons. Re-analysis of RNA-seq data from *C9ORF72* ALS/FTD post-mortem tissue^65^ confirmed this observation, demonstrating a >50% drop in *ASS1* levels selectively in neurons with TDP43 pathology (**Fig. 6d**). As *ASS1* loss is associated with neurological phenotypes in citrulinemia^60^, we asked whether *ASS1* loss-of-function is sufficient for neurotoxicity. For this, we utilized automated microscopy and survival analysis, a technique we optimized for monitoring the survival of rodent primary neurons in several different disease-associated contexts^11,12,61,66,67^. Mixed cortical neurons were isolated from non-transgenic rodents, then transfected with EGFP and an RFP-tagged plasmid encoding either shRNA targeting the coding sequence of *ASS1* or a scrambled sequence. Transfected neurons were followed daily via fluorescence microscopy, and individual neurons were identified by image segmentation software (**Fig. 6e**). Time of death for each cell was determined by morphological criteria (neurite retraction, soma rounding, dissolution or loss of fluorescence) that proved to be highly sensitive indicators of cell death^68^. Taking advantage of the variability of transient transfection in neurons^12,68,69^, we stratified neurons by RFP fluorescence (encoded on the same plasmid as *ASS1* shRNA), and selected cells expressing high or low levels of shRNA. Compared to scrambled shRNA, shASS1 high-expressing neurons displayed a significant reduction in ASS1 protein, whereas shASS1 low-expressing neurons exhibited a non-significant trend towards lower ASS1 levels (**Fig. 6f**). We then compared the survival of neurons with low- or high-expression of ASS1 shRNA to those transfected with non-targeting shRNA (**Fig. 6g**). For both low and high-expressing neurons, *ASS1* depletion resulted in a significant increase in the cumulative risk of death compared to controls, suggesting that even slight reductions in *ASS1* expression can be neurotoxic.

**Figure 6:**
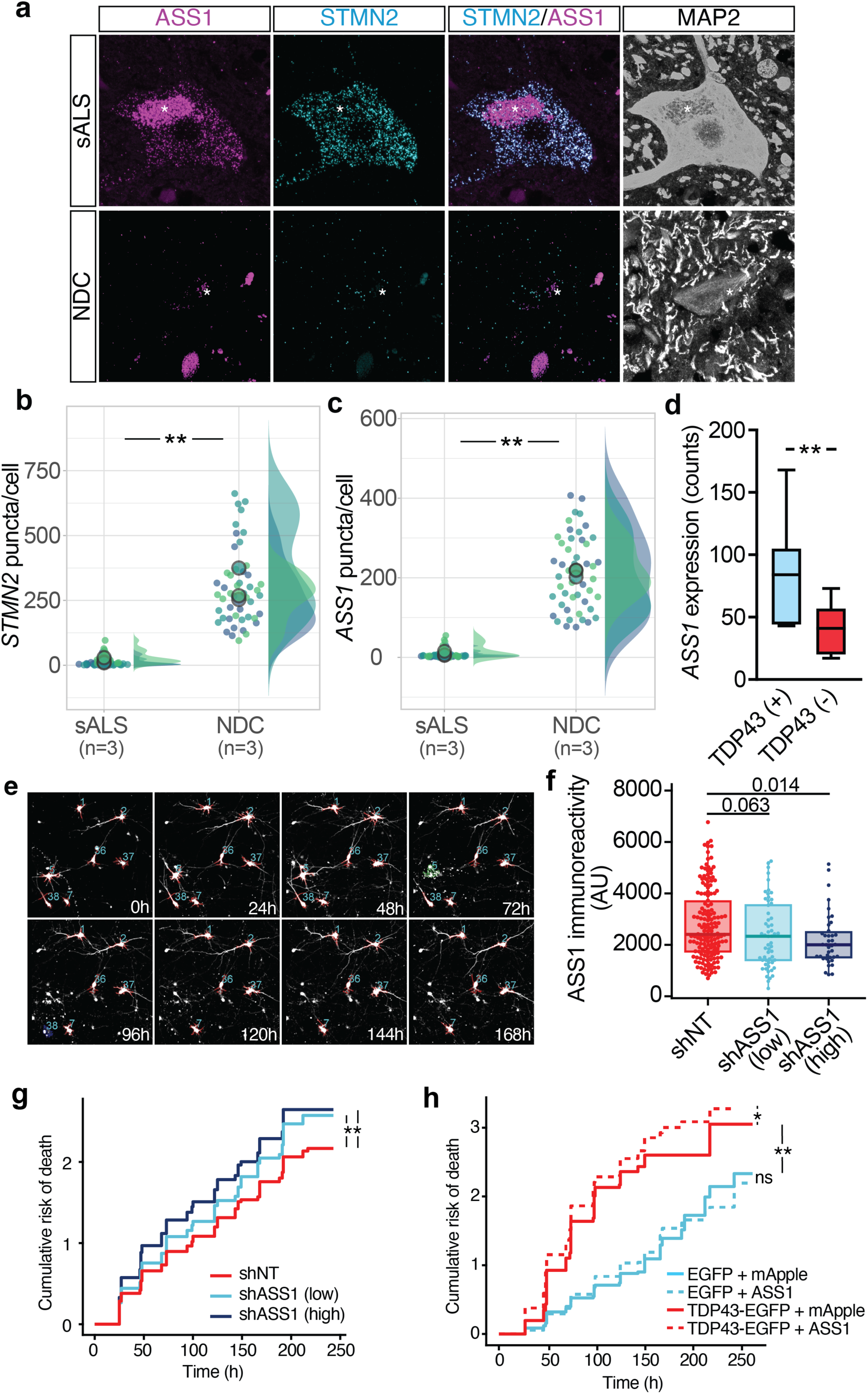
ASS1 is an essential factor reflecting TDP43 function in ALS. **a**, Single-molecule fluorescence in situ hybridization (smFISH) for *STMN2* and *ASS1* in post-mortem human ALS and neurological disease control (NDC) spinal cord. All tissues were immunostained for MAP2 to aid in identification of motor neurons within the anterior horn. Count of *STMN2* (**b**) and ASS1 (**c**) smFISH puncta in motor neurons from ALS and NDC spinal cord. **p<0.05, Kruskal-Wallis with Dunn’s post-hoc test. **d**, Abundance of ASS1 in neurons displaying TDP43 pathology, sorted from the brains of C9ORF72-ALS/FTD patients (adapted from Liu et al., 2019)^65^. **p_adj_=0.002, Wald test with Benjamini–Hochberg correction. **e**, longitudinal microscopy of rodent primary cortical neurons expressing mApple. Neurons are imaged repeatedly over a 10d period, identified by segmentation algorithms (red outlines) and assigned a unique identifier (blue numbers). Cell death is indicated by fragmentation and loss of neurites (green outline) and/or reduction in fluorescence intensity (dark blue outline). **f**, ASS1 immunoreactivity in neurons transfected with non-targeting (NT) or ASS1-directed shRNA, separated into low- and high-expressing groups based on shRNA reporter levels. Groups are compared by Wilcoxon rank-sum test. **g**, cumulative risk of death for primary neurons transfected with shASS1 or non-targeting shRNA. **h**, cumulative risk of death for primary neurons expressing EGFP or TDP43-EGFP, together with mApple or ASS1. In **g** and **h**, data combined and stratified among 4 separate experiments imaged every 24h for 10d (*p≤0.01, **p<2×10^-16^, Cox proportional hazards; n=733-2288 neurons/condition).

Conversely, we asked whether restoring *ASS1* levels could mitigate TDP43-mediated toxicity. We transfected primary neurons with plasmids encoding either EGFP or TDP43-EGFP, together with mApple or ASS1 tagged with mApple, and imaged daily for 10d as before (**Fig. 6h**). As in prior investigations^11,12,61,66,69^, TDP43-EGFP overexpression resulted in a significant increase in the cumulative risk of death compared to EGFP alone. Although ASS1 expression had no apparent effect on the survival of EGFP-expressing control neurons, ASS1 over-expression resulted in a modest, yet significant increase in risk of death for TDP43-EGFP-expressing neurons, suggesting that ASS1 restoration is insufficient to rescue TDP43 related toxicity, and in fact can compound the toxicity associated with TDP43 accumulation.

### UPF1 and TDP43 counter-regulate global 3’UTR length and RNA stability

Independent of its splicing activity, TDP43 can modulate RNA stability by regulating polyA usage and 3’UTR length^27,70,71^. We therefore wondered if changes in the expression of *ASS1* and other genes identified in TDP43 over-expressing iMNs correlate with 3’UTR usage. To do this, we applied DaPars^72^ to detect differential use of novel and annotated polyA sites in our lentivirus-treated iMN RNA-seq dataset. Using the mean percentage of <u>D</u>istal <u>p</u>oly<u>A</u> site <u>U</u>sage (DPAU), we found 4280 3’UTR-lengthening changes and 267 3’UTR-shortening events in TDP43-overexpressing iMNs, compared to iMNs transduced with EGFP alone (**Fig. 7a**). This is consistent with previous illustrations of proximal alternative polyA sites and shortened 3’UTRs upon TDP43 knockdown^26,27,71^. Comparing UPF1 and TDP43 co-expressing iMNs to EGFP expressing iMNs, we found only 536 3’UTR-lengthening and 787 3’UTR-shortening events (**Fig. 7b**), confirming the link between 3’UTR length and a UPF1-dependent, PTC-independent mRNA decay mechanism^34,73,74^. To explore whether similar changes were present *in vivo*, we applied DaPars to RNA-seq data from *C9ORF72*-ALS/FTD frontal cortex^75^. These analyses uncovered 335 3’UTR-lengthening and 57 3’UTR-shortening PAU events in *C9ORF72*-ALS/FTD cortex compared to that from healthy controls (**Fig. 7c**). Together these data highlight 3’UTR lengthening is a conserved feature of ALS/FTD pathology, and furthermore suggest that UPF1 expression can partially reverse the observed increases in distal polyA site usage.

**Figure 7:**
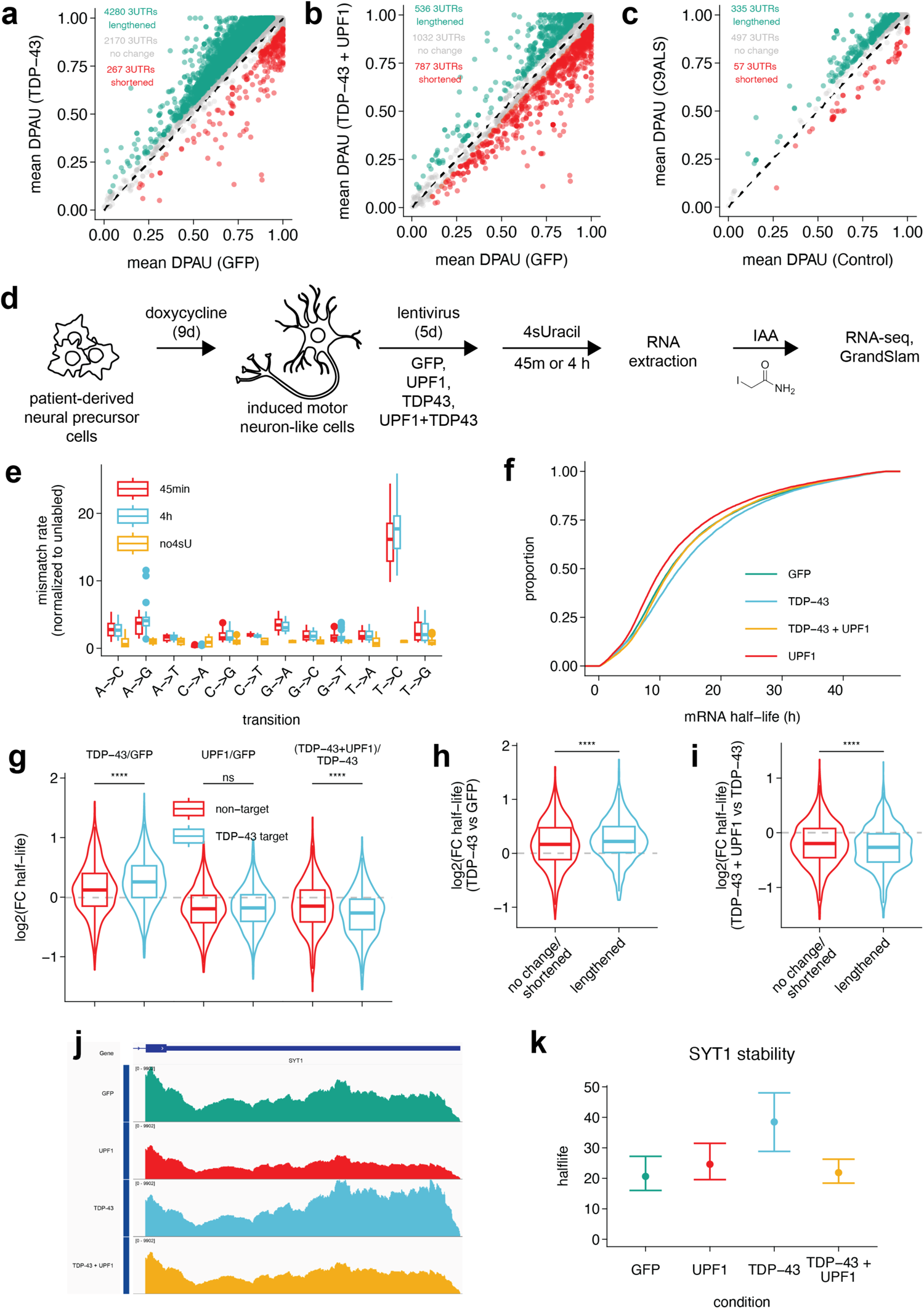
UPF1 restores global 3’UTR length and RNA stability in iMNs overexpressing TDP43. **a-c**, mean distal polyA site usage index (DPAU) scatter plots in (**a**) TDP43 vs GFP expressing iMNs, (**b**) TDP43/UPF1 vs GFP expressing iMNs, and (**c**) C9ALS vs Healthy control frontal cortex. **d**, experimental design of SLAM seq in lentivrus treated iMNs. Labeling and sequencing was conducted in biological triplicate. **e**, mismatch rate of each nucleobase as determined by deep RNA sequencing of 4sU-labeled and unlabeled iMNs. **f**, empirical distribution function of RNA half-lives in hours from iMNs expressing indicated lentivirus constructs. **g**, distribution of fold changes in RNA half-lives for indicated comparison (top) among TDP43 targets previously identified in CLIP-seq studies^81,82,106^. **h**-**i**, distribution of fold changes in RNA half-lives (TDP43 vs GFP in **h**, or TDP43/UPF1 vs TDP43 in **i**) for genes that exhibited significant 3’UTR lengthening or non-lengthening changes in TDP43-expressing iMNs compared to GFP-expressing iMNs. Data points in **g**-**i** represent gene-level estimations of RNA half-lives. Distributions are compared by Wilcoxon rank-sum test (****: p < 2 x 10-16). **j**-**k**, representative example of increased distal polyA site usage (**j**) and half-life in hours (**k**) of synaptotagmin 1 (*SYT1*). Error bars represent 95% confidence interval.

As TDP43 and UPF1 induced counter-directional changes in polyA site usage and 3’UTR length, and both factors are intricately connected with RNA stability, we asked whether RNA turnover is likewise affected oppositely by TDP43 and UPF1. To examine RNA clearance in iMNs, we took advantage of metabolic pulse-chase labeling of nascent RNA using thiol(SH)-containing uridine analogs, coupled with RNA-sequencing (SLAMseq)^76^. Lentivirus-treated iMNs were incubated with 4-thiouridine (4sU) for 45m to label actively-transcribed RNA, and then washed with unlabeled uridine (**Fig. 7d**). For each condition, samples were collected immediately after the labeling period as well as after 4h of uridine chase. Purified 4sU-containing RNA was chemically alkylated, leading to mis-interpretation of labeled uridine residues as cytosine by next-generation sequencing. Transitions are then quantified in each read to determine the approximate fraction of “new” and “old” RNA as well as half-life on a gene-by-gene basis using GRAND-SLAM^77^.

To determine the selectivity of labeling, we first examined the relative rate of transitions by nucleobase (**Fig. 7e**). In the majority of 4sU-labeled samples, we observed the expected T-to-C transitions at nearly 20-fold the rate of unlabeled iMNs, consistent with efficient labeling. The rate of off-target labeling, on the other hand, was less than 3-fold in the majority of cases. We next examined the effect of UPF1 and TDP43 overexpression on global RNA stability (**Fig. 7f**). UPF1 overexpression induced a leftward shift in the distribution of RNA half-lives compared to EGFP, consistent with broad RNA destabilization and reflective of the canonical function of UPF1 in myriad RNA decay pathways. TDP43 overexpression, on the other hand, induced a rightward shift in RNA half-lives, suggesting overall transcript stabilization. This conflicts with previous data showing a net destabilization of RNA in iPSCs transiently expressing TDP43^31^, potentially a result of cell type-specific consequences of TDP43-driven 3’UTR lengthening. That is, whereas 3’UTR shortening is generally stabilizing in most mammalian cell types^78,79^, 3’UTR lengthening may uniquely stabilize RNA in neurons^80^. Notably, coexpression of UPF1 and TDP43 yielded a distribution of RNA half-lives that was identical to that of EGFP overexpression, suggesting that UPF1 and TDP43 counter-regulate the stability of the neuronal transcriptome as a whole.

To further understand the determinants of UPF1- and TDP43-driven changes in RNA stability, we first categorized transcripts as direct TDP43 RNA targets or non-targets, based on available TDP43 CLIP-seq data^81,82^ (**Fig. 7g**). In doing so, we observed that TDP43 targets were preferentially stabilized by TDP43 overexpression in comparison to non-targets, suggesting a selective effect of TDP43 on RNA half-life driven by direct associations of TDP43 and its target transcripts. Conversely, the destabilizing effect of UPF1 was unselective, consistent with previous findings that the nature of UPF1 target discrimination is largely independent of sequence motifs^73,83,84^. We also asked whether genes demonstrating differential polyA site usage via steady-state RNA-seq were differentially stabilized by TDP43 expression as determined by SLAM-seq. Indeed, genes that exhibited TDP43-driven 3’UTR-lengthening were significantly more stabilized than those that exhibited no change in 3’UTR length or those that displayed 3’UTR-shortening (**Fig. 7h**). Additionally, genes that exhibited TDP43-driven 3’UTR-lengthening were significantly more destabilized by UPF1 coexpression compared to genes showing no change in 3’UTR length or 3’UTR-shortening (**Fig. 7i**). These data suggest that TDP43-induced 3’UTR-lengthening events primarily stabilize gene products, an effect that is reversed by UPF1 expression. For example, the gene encoding vesicle membrane protein synaptogamin (*SYT1*) exhibited 3’UTR-lengthening changes upon TDP43 overexpression and in *C9ORF72*-linked ALS/FTD frontal cortex (**Fig. 7j**), and increased stability in TDP43-overexpressing iMNs. UPF1 co-expression normalized *SYT1* stability in iMNs, however, in conjunction with restored 3’UTR length and polyA site usage (**Fig. 7k**). These data suggest that TDP43 preferentially drives distal polyA site usage, conferring increased stability, and that UPF1 coexpression eliminates 3’UTR-lengthened transcripts through a PTC-independent and 3’UTR length-dependent decay mechanism.

## Discussion

In this study, we explored whether NMD was impaired in ALS and UPF1-treated models of ALS/FTD. To accomplish this we devised a next generation sequencing-approach to quantify the abundance of PTC-containing transcripts in a platform-independent manner. After validating our approach with genetic and pharmacologic models of NMD impairment, we applied our pipeline to ten ALS/FTD RNA-seq datasets as well as one Huntington disease RNA-seq dataset. We found no evidence of elevated PTC-containing transcripts in disease contexts. In fact, several iMN models indicate subtle decreases in NMD-sensitive reads, suggesting subtle NMD enhancement. In the absence of obvious NMD impairment, we expanded our search for PTC-independent mechanisms of UPF1-related neuroprotection. We identified *ASS1* as a TDP43-downregulated and UPF1-normalized gene whose loss of function has critical implications for neuronal health, and whose expression is dramatically reduced in spinal neurons from ALS patients. Finally, we demonstrated that TDP43 and UPF1 have antagonistic effects on steady state 3’UTR length as well as RNA stability in a manner consistent with PTC-independent, long-3’UTR-triggered UPF1 activity. These findings challenge the prevailing PTC-based model of UPF1-mediated neuroprotection and provide the basis for an underappreciated role of UPF1 in selectively clearing transcripts with abnormally long 3’UTRs. Our observations are mirrored by concurrent studies from Alessandrini et al., confirming the accumulation of transcripts with long 3’UTRs in patient-derived samples and human neuron ALS models. Together, these investigations argue for a novel mechanism of neuroprotection by UPF1 involving the clearance of alternatively polyadenylated transcripts with long 3’UTRs induced by TDP43 dysfunction, a highly prevalent and signature neuropathological event in the majority of those with ALS and FTD.

To identify novel and broadly expressed NMD substrates in an unbiased manner, we implemented a bioinformatics pipeline and applied both global and candidate-based strategies to characterize NMD function in ALS/FTD models and disease. The pipeline, termed NMDq, was constructed from widely available software that, to our knowledge, had not previously been used as presented to explore the pathophysiology of ALS, FTD, or other neurological conditions. To validate the accuracy and biological relevance of NMDq, we first analyzed 6 RNA-seq datasets of NMD-impaired cell systems: 3 from publicly available repositories, 2 from collaborators, and 1 from this study. We also included data from a variety of cell types with both genetically and pharmacologically-driven NMD impairment, ensuring the universality of our approach. We devised two complimentary statistics that effectively summarize NMD activity from sequencing data: global NMD burden (GNB), or the cumulative TPM of transcripts containing PTCs, and differential NMD-class usage (DNU), or the relative preference for PTC-containing transcripts on a gene-by-gene basis. Using GNB, we can draw conclusions about the global capacity of represented cell systems to surveil and metabolize derangements in the transcriptome pertaining specifically to canonical NMD affecting PTC-containing transcripts. In contrast, using DNU we can assay the sensitivities of specific genes to the inhibition of NMD by way of the balance of NMD-sensitive to NMD-insensitive products.

Taking this further, we used a set analysis of DNU-positive genes (genes that showed significant differential NMD-class usage) across all included datasets to identify consistent and reproducible PTC-containing NMD substrates. We used the isoform expression characteristics of NMD-sensitive gene products to identify eight candidates for which we designed isoform-specific and total gene qRT-PCR primers. Several of these candidate substrates, such as *SRSF2*, *SRSF3*, *SRSF6*, and *TRA2B,* belong to a family of serine-arginine splicing factors that autoregulate themselves by directing the inclusion of a PTC-containing “poison” exon^45^ (as in **Supp. Fig. 1c**). Other targets, including *BAG1* and *SAT1*, are subject to NMD due to alternative splicing and the inclusion of conserved exons carrying stop codons^48,85^. Several substrates, including *TRIR* and *DDX1*, have no prior association with NMD and are therefore novel targets; the latter of these (*DDX1*), is a neuronal-selective marker of NMD that may be invaluable for future studies of cell type-specific NMD regulation in neurodegenerative disease models.

To validate our candidate NMD-substrates, we implemented both steady-state and nascent qRT-PCR. At steady-state, all candidates showed significantly improved sensitivity to pharmacologic NMD inhibition compared to previously published targets, including *ATF3*, *ATF4*, and *ERN1*^47^. To explore whether these changes were due to a post-transcriptional destabilization event or an increase in transcription, we employed roadblock-qPCR^49^, a significantly less toxic and more direct approach to studying RNA stability compared to the most common approach involving transcriptional inhibitors such as actinomycin-D^86^. Our results show that NMD inhibition results in upregulation of *BAG1*, *SAT1*, and *SRSF2* primarily via stabilization of otherwise highly unstable transcripts. Importantly, enhancements in transcript stability were also associated with modest increases at the protein level for SAT1. Additional studies are warranted to determine the stability of the resulting truncated protein product and the extent to which antibody or epitope selection influences the detection of such shortened or out-of-frame products.

Encouraged by the ability of NMDq to accurately identify changes in NMD activity at the global level as well as for individual transcripts and genes, we applied the pipeline to ten ALS/FTD RNA-seq datasets and one from Huntington’s disease. We did not find convincing evidence of NMD dysfunction in either cell-based models or human post-mortem tissue, challenging the prevailing suspicion that UPF1 overexpression prevents neurodegeneration by restoring or enhancing canonical NMD activity. Although growing evidence points towards altered RNA metabolism in ALS models and disease^31,87,88^, our data suggest that such impairments are not a consequence of insufficient capacity to degrade or otherwise resolve PTC-containing transcripts. In fact, in datasets relevant to *C9ORF72*-related ALS/FTD as well as sporadic ALS, we noted subtly *<u>reduced</u>* NMD-substrate burden consistent with *<u>increased</u>* canonical NMD. A notable exception to this global trend of intact NMD function may be the preponderance of NMD-substrates in HEK293 cells overexpressing FUS or mutant variants of FUS. Consistent with previous studies^88^, we found both WT and disease-associated FUS mutations elicit small increases in NMD-sensitive gene products of *BAG1*, *SAT1*, and *SRSF2*. We also failed to find any evidence of differentially active, phosphorylated UPF1 in post-mortem tissue from ALS patients. Taken together, these observations argue against global NMD impairment in ALS.

Without clear evidence of NMD dysfunction, specifically with regard to the clearance of PTC-containing transcripts, we expanded our search for mediators of UPF1-associated neuroprotection through conventional, gene-based analyses of RNA-sequencing data from iMNs overexpressing TDP43 and/or UPF1. In the process, we identified several genes that responded robustly and antagonistically to TDP43 and UPF1 overexpression. One such gene, *ASS1*, encodes an essential enzyme that catalyzes the rate-limiting step in arginine biosynthesis. Loss of ASS1 function results in type 1 citrullinemia, a primary metabolic disorder characterized by significant neurologic impairment^60^. Our studies show that TDP43, but not RNA binding-deficient TDP43 (F147L, F149L), drives the downregulation of this vital gene, suggesting that this effect is mediated via direct interaction of TDP43 with the *ASS1* transcript. Biotinylated ASO pull-down assays confirmed this possibility, revealing direct interactions between *ASS1* RNA and both TDP43 as well as UPF1. In keeping with the TDP43-dependent downregulation of *ASS1* we observed in iMNs, we also detected a dramatic reduction in *ASS1* expression within spinal motor neurons of ALS patients, compared to controls. *ASS1* was also significantly downregulated in neurons with TDP43 pathology, as detected by prior studies involving fluorescence-based sorting of neuronal nuclei from *C9ORF72* ALS/FTD post-mortem cortex^65^. Furthermore, longitudinal microscopy and automated survival analysis of primary neurons confirmed the importance of *ASS1* in preventing neuron loss, in accord with previous studies showing that neurons are highly sensitive to both decreased and increased ASS1^60,89,90^. ASS1 overexpression was insufficient to rescue TDP43 toxicity, however. There are at least two explanations for this finding. First, TDP43 regulates thousands of transcripts^81,82^, many of which including *ASS1*, are crucial. Therapeutic restoration of *ASS1* in and of itself may not be sufficient to prevent TDP43-driven toxicity. Second, *ASS1* restoration may require finer control of gene expression than is possible with transient transfection. In murine models of citrullinemia, for example, *ASS1* gene therapy using adeno-associated virus prevents premature death and lowers plasma citrulline, but paradoxically results in arginine depletion and renal failure through unknown mechanisms^90^.

To further elucidate the means by which TDP43 and UPF1 counter-regulate gene expression, we quantified differential polyA site usage using DaPars^72^. Prior studies suggested that TDP43 binds to and influences the use of distal polyA sites on nascent RNA, thereby affecting the expression of TDP43 itself^71^ as well as many other genes^70^. In accord with these findings, our data demonstrate preferential distal polyA site usage in TDP43-overexpressing iMNs, together with an overall increase in global RNA stability. UPF1, on the other hand, has no established influence on polyA site usage. Instead, it may regulate mature mRNA levels by sensing 3’UTR length and potentiating EJC- and PTC-independent transcript decay^73,84,91–93^. Co-expression of UPF1 prevented TDP43-induced distal polyA site usage, and effectively countered changes in RNA stabilization associated with TDP43 expression. Although additional studies are required to directly interrogate the mechanisms behind UPF1-mediated length-sensing of TDP43-lengthened transcripts, these investigations are in keeping with the consistent ability of UPF1 expression to prevent TDP43-related toxicity in multiple model systems, from yeast to primary and human neurons, flies, and rodents^12,13,19–22^.

We also asked whether the changes in polyA site usage associated with TDP43 and UPF1 expression might correlate with changes in global RNA stability. To answer this question, and to identify candidate mediators of UPF1-driven rescue that were not evident from steady-state RNA-seq studies, we employed SLAM-seq^76,77^, wherein total RNA is sequenced and estimations of nascent RNA synthesis and stability are derived from C-T transitions induced by 4-thiol-uridine labeling. We found that global RNA stability was reduced by the expression of UPF1 and increased by the expression of TDP43. The observed effect of UPF1 expression on RNA stability is consistent with all canonical and non-canonical functions of UPF1 as an RNA decay factor. TDP43’s effects on RNA stability, however, conflict with previous findings in undifferentiated iPSCs, obtained through the related technique BruChase-seq^31^. This discrepancy could be due to a number of factors. First, determinants of RNA stability are cell type-specific^43,94^. A relevant example of this is the differential impact of 3’UTR length on RNA stability in neurons compared to non-neuronal cells. In neurons, 3’UTR lengthening stabilizes transcripts, whereas the opposite relationship between 3’UTR length and RNA stability is observed in non-neuronal cell types^80,95,96^. Second, differences in the techniques used to study RNA stability may account for some of the observed experimental discordance. In BruChase-seq, for example, bromouridine-containing RNA is purified from total RNA via immunoprecipitation using anti-bromouridine antibodies prior to sequencing^97^. SLAM-seq avoids the prohibitively high input requirements for BruChase-seq. The consequences of these technical distinctions have not been directly interrogated, but may stem from differences in cDNA library depth and coverage of modified RNA. In SLAM-seq, transition-containing reads represent only a small fraction of total reads which may result in sample error. In contrast, all reads are bromouridine-containing in BruChase-seq, but unmodified reads are unavailable to provide additional insight into differences in steady-state RNA abundance that precede labeling. Together, these emerging techniques offer complimentary perspectives on our developing understanding of RNA stability in experimental cell systems.

## Materials and methods

### NMDq

Paired-end RNA-seq reads from publicly accessible datasets were quality and adapter trimmed (TrimGalore v0.6.6, default parameters) and aligned to the human genome (STAR v2.7.3a; GRCh38, Gencode v35) with the optional parameters: “--twopassMode Basic --outSAMtype BAM SortedByCoordinate --outSAMattributes NH HI NM MD AS nM jM jI XS --quantMode GeneCounts”^35^. For each dataset, a de novo transcriptome inclusive of all identified junctions was assembled (“--merge”) and BAM files were requantified (StringTie v2.1.4, default parameters)^36,37^. TPMs were calculated and open reading frames (ORFs) were estimated to predict premature termination codons (PTCs) (IsoformSwitchAnalyzeR v1.10.0)^38^. Transcripts were categorized into “NMD” or “non-NMD” classes according to their Ensembl annotated or predicted sensitivity to NMD. TPMs were aggregated class- and gene-wise and summarized with two approaches: 1) Global NMD burden: Group-wise sums of NMD-TPM was as a proportion of total TPM was and normalized to within-dataset controls. 2) Differential NMD-class usage (DNU): Gene-wise ratios of NMD to total transcript TPM were calculated (NMD ratio). NMD ratio was log-transformed and used to fit generalized linear models to gene- and group-wise comparisons (limma v3.44.3)^98^. An empirical Bayes test was used on the resulting model to determine statistical significance. Genes showing ≥1.5 fold change from compared conditions were counted as exhibiting DNU.

### HEK293T cell culture

Human embryonic kidney (HEK) 293T cells were cultured in DMEM (GIBCO), 10% FBS, 100 units/mL Penicillin/Streptomycin at 37°C in 5% CO2. Cells were passaged when they reached 75-80% confluency using Accutase (ThermoFisher #MT25058CI), and transfected using Lipofectamine 2000 (Fisher Scientific #11668019) according to the manufacturer’s protocol.

### iMotor neuron differentiation and transduction

iMotor neurons were differentiated as described in Chua et al. (2022)^99^. Briefly, iMotor neuron neuroprogenitors were thawed in TeSR-E8 (Stemcell Technologies) containing 2ug/ml doxycycline (Sigma #D3447) and ROCK inhibitor (Fisher Scientific #BDB562822) on plates coated with matrigel (Fisher Scientific #354230) or polyornithine (Sigma #P3655) and laminin (Fisher Scientific #CB40232). The following day, the media was changed to TeSR-E8 containing 2ug/ml doxycycline, 1x N2 supplement (Gibco, #17502-048), 1:10,000 Compound E (Millipore #565790; day 1). On day 3, the media was changed to DMEM/F12, 1X N2 supplement, 1x NEAA Supplement (Gibco, #11140-050), 1x Glutamax Supplement (Gibco, #35050-061), 2ug/mL doxycycline, 1:10,000 Compound E. On day 6, the media was changed to Neurobasal-A (Gibco, #12349-015), 1x B27 supplement (Gibco, #17504-044), 1x Glutamax, 10 ng/mL BDNF (Peprotech, #450-02), 10 ng/mL NT3 (Peprotech, #450-03), 0.2 μg/mL laminin (Sigma, #L2020), 2ug/mL doxycycline, 1:100 Culture One (ThermoFisher #A33202-01). Depending on the length of the experiment, additional day 6 media (excluding Culture 1) was added every 3 days. On day 9, cells were transduced with the appropriate virus (prepared by University of Michigan Vector Core) and cells were sustained in the same culture medium for the remainder of the experiment. On day 11, iMNs were treated with 1uM 11j SMG1-inhibitor or an equivalent volume of DMSO. iMNs were collected for RNA isolation on day 14.

### Next-generation sequencing

cDNA libraries were prepared from Trizol extracted, DNA-digested samples using the Illumina Stranded mRNA prep kit (Illumina, #20040532). Paired end sequencing was carried out on an Illumina NovaSeq (S4) 300 cycle sequencer at the University of Michigan Advanced Genomics Core. Samples were sequenced at an average depth of 75M reads/sample.

### Roadblock qPCR

Live HEK293T or iMN cells were metabolically labeled for 6h with the nucleoside analog 4-thiouridine (4sU, final concentration 400uM) in 2h intervals. Following RNA extraction and DNase treatment, incorporated 4sU was modified using the thiol-reactive compound N-ethylmaleimide (NEM, 50ug/ul) in a compatible buffer (125mM Tris-HCl pH 8.0, 2.5mM EDTA, ultrapure water). 0.2ng/ul firefly luciferase mRNA (FL, synthesized in house) was spiked into thiol modified samples. Reverse transcription was performed using the ProtoScript® II First Strand cDNA Synthesis Kit (NEB, #E6560S) and 18-mer oligo dT primers. Quantitative PCR was carried out using PowerUp SYBR Green Master Mix (ThermoFisher #A25742). Target CT values were normalized to the geometric mean of GAPDH and FL.

### Primary neuron cell culture and transfection

Cortices from embryonic day (E)19–20 Long-Evans rat embryos were dissected and disassociated, and primary neurons plated at a density of 6×10^5^ cells/mL in 96-well plates, as described previously^100^. At in vitro day (DIV) 4–5, neurons were transfected with 100 ng of pGW1-mApple^69^ to mark cells bodies and 100 ng of an experimental construct (i.e. FUGW-TDP43-EGFP) using Lipofectamine 2000, as before^100^. Following transfection, cells were placed into Neurobasal with B27 supplement (Gibco, Waltham, MA; for all survival experiments). For shRNA knockdown experiments, neurons were transfected with 50 ng of FUGW-EGFP and 200ng shRNA/well. Cells were treated with either SMARTvector Lentiviral Rat control hEF1a-TurboRFP shRNA (Horizon, Lafayette, CO) or SMARTvector Lentiviral Rat Ass1 hEF1a-TurboRFP shRNA (5’-ATTCCAATGAAGCGGTTCT-3’) targeting a coding region in the 12th exon of rat ASS1 (V3SR11242-238358331).

### Longitudinal fluorescence microscopy

Neurons were imaged as described previously^11,12,61,66,67,69^ using a Nikon (Tokyo, Japan) Eclipse Ti inverted microscope with PerfectFocus3 and a 20X objective lens. Detection was accomplished with an Andor (Belfast, UK) iXon3 897 EMCCD camera or Andor Zyla4.2 (+) sCMOS camera. A Lambda XL Xenon lamp (Sutter) with 5 mm liquid light guide (Sutter Instrument, Novato, CA) was used to illuminate samples, and custom scripts written in Beanshell for use in μManager controlled all stage movements, shutters, and filters. Custom ImageJ/Fiji macros and Python scripts (https://github.com/barmadaslab/survival-analysis and https://github.com/barmadaslab/measurements; copies archived at https://github.com/elifesciences-publications/survival-analysis and https://github.com/elifesciences-publications/measurements) were used to identify neurons and draw regions of interest (ROIs) based upon size, morphology, and fluorescence intensity. Criteria for marking cell death included rounding of the soma, loss of fluorescence and degeneration of neuritic processes.

### Immunohistochemistry

Immunostaining was accomplished using the Dako Autostainer Link 48 (Agilent, USA). An anti-pUPF1 antibody (Milipore #07-1016, 1:300) was used with the Dako High pH Target Retrieval Solution (Tris/EDTA, pH 9; Agilent, USA) (20 minutes, 97°C) and the Dako Envision Flex Plus Mouse Link Kit (Agilent, USA) to detect the antibody along with the Dako DAB (Agilent, USA). Whole-slide images were generated by the University of Michigan Digital Pathology group within the Department of Pathology using a Leica Biosystems Aperio AT2 scanner equipped with a 0.75 NA Plan Apo 20x objective; 40x scanning is achieved using a 2x optical magnification changer. Resolution is 0.25µ/pixel for 40x scans. Focus during the scan is maintained using a triangulated focus map built from individual focus points determined in a separate step before scanning is started. Proprietary software is used for image processing during acquisition. Image analysis performed using QuPath software and mean intensity calculated for stained cells.

### ASS1 mRNP pull down assay

Purification of target-specific mRNPs was adapted from Matia-González et al. (2017)^64^. Briefly, HEK293T cells grown on standard 10cm culture dishes were UV-crosslinked with 254nm UV at 150 mJ/cm2. Cells were harvested by triturating and pelleted at 235g for 10m, discarding the supernatant. The cell pellet was resuspended in 200µl of ice-cold lysis buffer (100 mM Tris–HCl pH 7.4, 500 mM LiCl, 10 mM EDTA, 1% Triton X-100, 5 mM DTT, 20 U/ml Baseline-ZERO DNase (Lucigen #DB0715K), 100 U/ml RNasin (Promega #N2611), and 1x cOmplete EDTA-free Protease Inhibitor Cocktail (Roche #11873580001)) and sonicated with 3 rounds of 20s bursts of level 10 amplitude Q55 Sonicator (Qsonica, Newtown, CT). The lysate was centrifuged at 15,000xg for 10m and the supernatant was collected. Roughly 35ug of protein and 100ul B&W buffer (10 mM Tris-HCl, pH 7.5, 150 mM NaCl, 0.5 mM EDTA, 32 U/ml RNasin, 10 U/ml Baseline-ZERO DNase, pH 8.0). Biotinylated ASOs were added to lysates to a final concentration of 22.50µM with respect to the volume of Dynabeads MyOne Streptavidin C1 slurry (Thermofisher #65001). The lysate-ASO solution was incubated at 70C for 5m in a thermoblock then removed and allowed to cool to room temperature. In a separate tube, 100µl of streptavidin beads/10cm plate was washed with an equal volume of B&W 3x, using a magnet to separate beads from the supernatant. Washed beads were added to the RT lysate-ASO solution and incubated for 30m at RT on a benchtop rotator. Beads were then collected via magnet and washed 3x with 500µl B&W warmed to 50C. After the final wash, beads were resuspended in 20µl Laemmli buffer for resolution with SDS-PAGE and western blotting.

### SLAM-seq

On the 14th day in vitro, live lenti-transduced iMNs were metabolically labeled with 400µM 4sU for 45m after which half were collected with Trizol and half were washed, incubated with 10mM unlabeled uridine for 4h, and collected with Trizol. After purification and DNase treatment, RNA was alkylated with a buffered iodoacetemide solution (10 mM iodoacetamide, 50% DMSO, 50 mM sodium phosphate buffer, pH8) at 50C for 15m. The alkylation reaction was stopped with 20mM DTT and the RNA was purified with ethanol precipitation. cDNA libraries were made using the Illumina Stranded Total RNA Prep with Ribo-Zero Plus kit (Illumina, #20040525). Paired end sequencing was carried out on an Illumina NovaSeq (S4) 300 cycle sequencer at the University of Michigan Advanced Genomics Core. Samples were sequenced at an average depth of 75M reads/sample. Trimmed reads were aligned to the hg38 genome using STAR (version 2.7.10b) with optional arguments “--outSAMtype BAM SortedByCoordinate --outSAMattributes NH HI NM MD AS nM jM jI XS --seedSearchLmax 20”^76^. Conversion analysis and kinetic modeling was achieved with GRAND-SLAM software and further analyzed in R with the GrandR package^77^.

### Fluorescence in-situ hybridization

In situ hybridization was performed per manufacturer’s recommended protocol (ACDbio Multiplex RNAscope). Fresh-frozen, paraffin-embedded post-mortem tissues mounted on slides were obtained from the University of Michigan Brain Bank (Table 3). Samples were de-paraffinized by immersion in xylene for 5m, then xylene for 5m with agitation, followed by 100% EtOH for 5m and 100% EtOH for 2m with agitation. The slides were dried for 5m at 60C before adding 5-8 drops of H_2_O_2_ (ACDbio #322381) to each and incubating at RT for 10m. Samples were rinsed with ddH_2_O 3-5 times before progression to antigen retrieval. Slides were placed in heated ddH_2_O (99C) for 10s, 1X retrieval agent (ACDbio #322001) for 15m at 99C, ddH_2_O for 15s at RT, 100%EtOH for 3m at RT, then dried at 60C for 5m. Outlines around each sample were drawn with a hydrophobic barrier pen (ACDbio #PN 310018), then 5 drops of Protease Plus (ACDbio #322330) were added prior to placing in a pre-warmed oven (40C) for 15m. Probes for STMN2 (ACDbio #525211-C2) and ASS1 (ACDbio #431291) were resuspended to 1X in probe diluent (ACDbio #300041) before baking in HybEZ II oven (ACDbio #321710/321720) for 2hr at 40C, then rinsing with 1X wash buffer (ACDbio #310091) for 2m at RT. Hybridization was accomplished with Multiplex FL v2 AMP1, AMP2 and AMP3 (ACDbio #323110), each at 40C for 30m, separated by rinses in 1X wash buffer for 2m at RT. For signal development, Multiplex FL v2 HRP-C1 was added to the samples for 15m at 40C. Slides were rinsed twice in 1X wash buffer for 2m at RT before adding 1X TSA Vivid Fluorophore 520 (ACDbio, #323271) for 30m at 40C, rinsing twice in 1X wash buffer for 2m at RT, then baking for 15m at 40C in Multiplex FL vs HRP Blocker (ACDbio, #323110) before rinsing 2x more in 1X wash buffer at RT. The process was repeated for Multiplex FL v2 HRP-C2 and -C3, using 1X TSA Vivid Fluorophore 570 and 1X TSA Vivid Fluorophore 650, respectively. Immunostaining for MAP2 was performed using chicken anti-MAP2 primary antibodies (Novus #NB300-213) diluted 1:3000 in 1% PBS with BSA, ON at 4C. Slides were rinsed three times in 1X wash buffer for 2m at RT before adding secondary antibody (goat anti-chicken Alexa 647, 1:250 in PBS with 1% BSA; Jackson Immunoresearch #103-605-155). Slides were rinsed five times in 1X wash buffer for 2m at RT, and coverslips were mounted using Prolong Gold (without DAPI; ThermoFisher #P36935). Full-thickness Z-stacks were acquired from the anterior horn of each slide using a Nikon (Tokyo, Japan) NSPARC confocal microscope and NIS Elements software. Puncta were quantified from maximal intensity projections of each image in ImageJ/FIJI by drawing regions of interest around individual neurons in the MAP2 channel, and counting puncta in remaining channels using the ‘analyze particles’ feature. Plots and statistical comparisons for puncta number/cell were created using Superplots and R, respectively^101^.

**Table 3:** Information on post-mortem samples.

| Sample | Age | Sex | PMI | Condition |
| --- | --- | --- | --- | --- |
| ALS-1 | 64 | Male | 7 | sALS |
| NDC-1 | 68 | Male | 22 | Alzheimer's disease |
| ALS-2 | 68 | Male | 4 | sALS |
| NDC-2 | 91 | Male | 18 | Alzheimer's disease |
| ALS-3 | 63 | Female | 7 | sALS |
| NDC-3 | 75 | Male | 6 | Alzheimer's disease |

NDC = non-ALS disease control

## Acknowledgements

We thank the patients that donated tissue samples to make this work possible. Skin samples from the study participants were collected and de-identified in collaboration with the Michigan Institute for Clinical and Health Research (MICHR, UL1TR000433) through an institutional review board (IRB)-approved protocol (HUM00028826). We thank Dr. Stephen A. Goutman, Director of the University of Michigan ALS Clinic and Biorepository, and Crystal Pacut from the Program for Neurology Research and Discovery. We also would like to acknowledge Mr. Matthew D. Perkins for his help with postmortem tissue from the University of Michigan Brain Bank, Kathy Toy for her expertise with immunohistochemistry, and Yi-Ju (LuLu) Tseng for her help in preparing RNA. Lastly, we are grateful to Francesco Alessandrini and Evangelos Kiskinis from Northwestern University for the generous use of RNA sequencing data.

This work was supported by National Institutes of Health (R01NS097542, R01NS113943, and 1R56NS128110-01 to S.J.B; F31NS115257 to N.G.; F31NS134123 to M.D.; P30AG072931 to the University of Michigan Brain Bank and Alzheimer’s Disease Research Center; T32-GM145470 to C.H. and the University of Michigan Cellular and Molecular Biology Graduate Program), the family of Angela Dobson and Lyndon Welch, the A. Alfred Taubman Medical Research Institute, the Danto Family, Ann Arbor Active Against ALS, and the Robert Packard Center for ALS Research. Immunohistochemistry was performed at the Rogel Cancer Center Tissue and Molecular Pathology Shared Resource Laboratory at the University of Michigan (NIH P30 CA04659229).

## Author contributions

N.G. and S.J.B. designed the study. N.G. performed all RNA isolations, bioinformatic analyses, transfections, wide-field fluorescence microscopy, RNA pull-downs and immunoblotting. F.A., E.K., B.Z., J.R., and J.K. provided RNA-seq data. C.A. and Y.B. performed fluorescence in-situ hybridizations and confocal microscopy. E.M.T. maintained iPSCs and differentiated neurons for experiments. X.L. assisted with cell culture, preparation of primary neurons, and RT-PCR. J.W. wrote the code for image acquisition and neuronal survival analysis. C.H. performed roadblock-qPCR. J.K., J.R. and B.Z. provided unique RNA-seq datasets; S.J.B. and N.G. assembled figures, wrote and edited the manuscript.

