## Supplemental material for "Counter-regulation of RNA stability by UPF1 and TDP43"

Sami Barmada

University of Michigan

Department of Neurology

109 Zina Pitcher Place, 4019 BSRB

Ann Arbor, MI 48109

**Supplemental Figure 1: NMDq identifies assayable, biologically relevant targets of canonical NMD (continued from Figure 3).**

**a**, First quadrant MA-plot (log-ratio vs mean expression) of iMN validation datasets (from left to right: Alessandrini, 2021: siUPF1-treated vs siControl-treated iMNs; Zaepfel, 2021: SMG1i-treated vs DMSO-treated iMNs; and Gomez, 2021: SMG1i-treated vs DMSO-treated iMNs). Points represent isoforms. Black points represent isoforms belonging to parental genes with at least one PTC-containing isoform up-regulated in two NMD-deficient conditions. **b**, SDS-PAGE and immunoblotting of HEK293T cells treated with DMSO or SMG1i and stained for GAPDH and SAT1. Data represents three biological replicates compared by paired Wilcoxon signed-rank test ($p = 0.25$). **c**, representative RNA-seq reads from DMSO- or 11j-treated iMNs of poison or putative poison exons at the *SRSF2*, *SAT1*, and *BAG1* loci.

**Supplemental Figure 2: NMD is not impaired in ALS models or tissues (continued from Figure 4).**

Cumulative TPM belonging to the PTC category in lentivirus-treated iMNs, normalized to GFP/DMSO treated condition. Data points represent biological replicates. Error bars represent standard error of the mean. Groups compared by Wilcoxon rank-sum test (*: $p<=0.05$).

### Supplemental figure 1

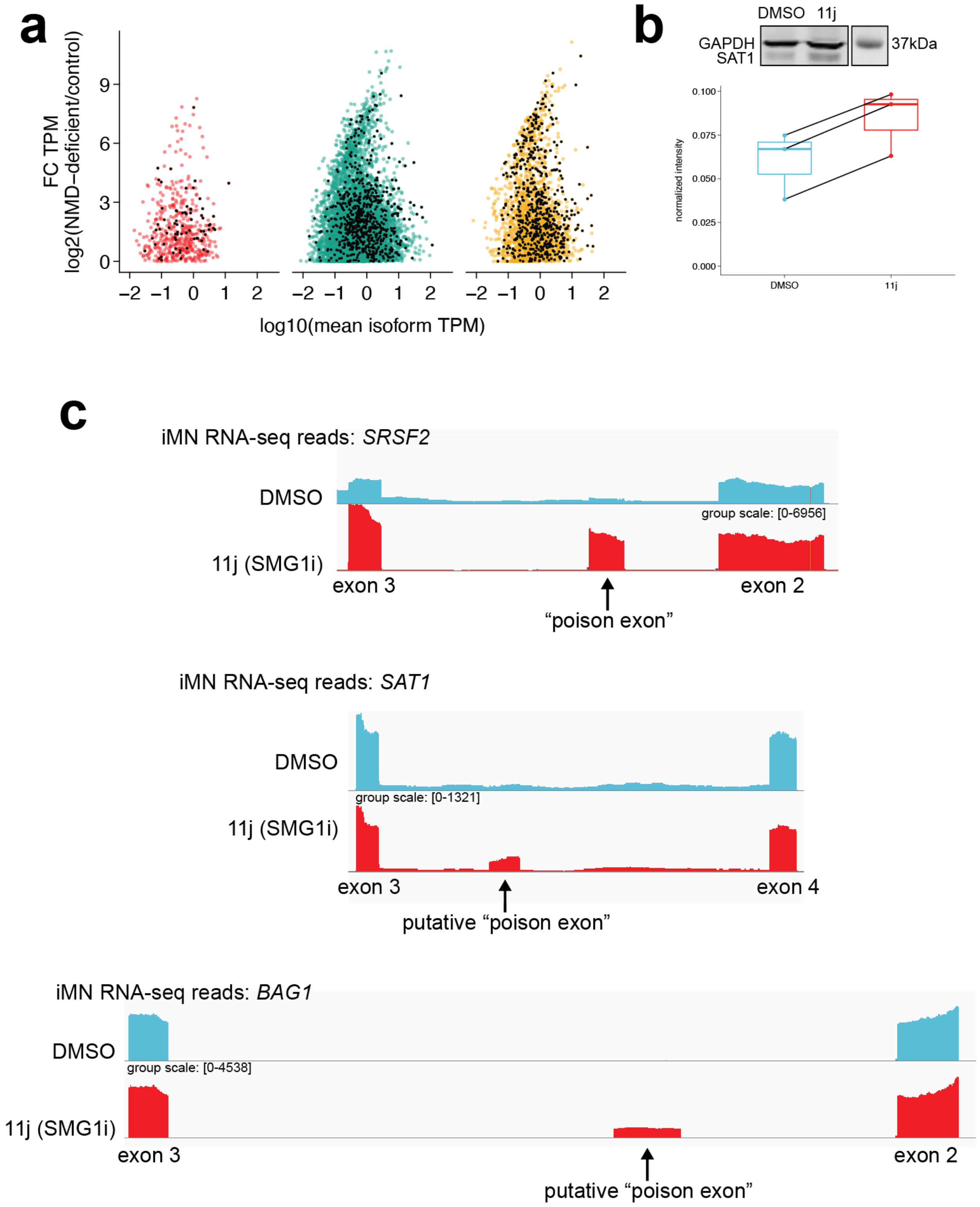

### Supplemental figure 2

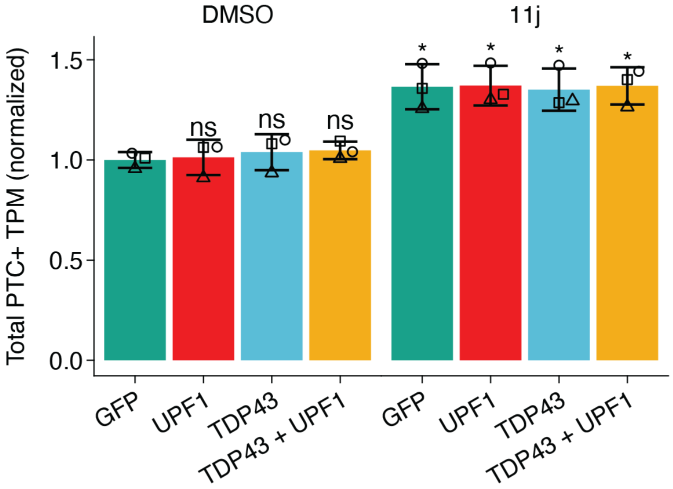
